# A human *GBA-L444P* transgene drives early and persistent dopamine neurotransmission deficits and alpha-synuclein pathology in a mouse model of early Parkinson’s disease

**DOI:** 10.64898/2026.05.25.727583

**Authors:** Natalie Connor-Robson, Tara Diviney, Javier Alegre-Abarrategui, Bradley Roberts, Katherine R Brimblecombe, Nora Bengoa-Vergniory, Harrison Waters, Milena Cioroch, Benjamin Davies, Kateryna O Bila, Martijn J C van der Lienden, Johannes M F G Aerts, Stephanie Cragg, Richard Wade-Martins

**Author notes:** Joint first authorship (Natalie Connor-Robson, Tara Diviney).

## Abstract

**Background:** Heterozygous mutations in the *GBA1* gene encoding the enzyme glucocerebrosidase (GCase) represent the most common genetic risk factor for developing Parkinson’s disease (PD). The underlying mechanisms by which *GBA1* mutations lead to PD through both loss- and gain-of-function effects remain unclear. There is a strong rationale for the generation and characterisation of a humanised *GBA1* mouse model to allow the effect of *GBA1* mutations on GCase function to be studied within the context of the human protein.

**Methods:** We have generated novel humanised mutant *GBA-L444P* and wild type *GBA-WT* mouse models using BAC recombineering and site-specific integration allowing the incorporation of the whole *GBA1* locus as a transgene, including the endogenous promoter, all exons and introns, and flanking regions. Our experimental design crossed each *GBA1* transgene onto a *Gba*^+/-^ background and included *Gba*^+/-^ littermate controls in our cohorts, allowing us to explore both the loss- and gain-of-function of *GBA1* mutations. We have carried out “deep phenotyping” to characterise these mice by biochemical, stereological and behavioural testing, and assess dopamine release and content using fast-scan cyclic voltammetry and high performance liquid chromatography.

**Results:** The *GBA-L444P* mice showed a significant reduction in GCase activity by 18 months of age and preferentially expressed a high molecular weight form of the GCase protein, likely due to retention in the ER and aberrant glycosylation. The *GBA-L444P,* but not *Gba*^+/-^, mice demonstrated an early and persistent reduction in dorsal striatal dopamine release in the absence of any dopaminergic cell loss or deficits in dopamine synthesis or reuptake, compared to human wild-type controls. *GBA-L444P* and *Gba*^+/-^ mice developed an accumulation of oligomeric α-synuclein pathology, but only *GBA-L444P* mice demonstrated subtle but significant changes in behaviour.

**Conclusions:** The novel humanised *GBA-L444P* mouse model described here helps to resolve gain- or loss-of-function effects of *GBA1* mutations seen in Parkinson’s as well as providing a novel set of models to investigate the human protein. Our work demonstrates that changes in dopamine release and behavioural deficits arise from a gain-of-function mechanism, whereas α-synuclein pathology arises from GCase loss-of-function.

## Introduction

Parkinson’s disease (PD) is characterised by the preferential degeneration of dopaminergic (DA) neurons in the substantia nigra pars compacta (SNpc) and by Lewy pathology [1]. PD is the second most common neurodegenerative disorder in the world, affecting approximately 1% of the global population over 60 [2]. Although 5-10% of PD cases directly arise due to Mendelian inheritance of a disease-causing genetic mutation [3], the majority of cases are attributed to a complex interplay of genetic and environmental risk-factors [4]. Aside from the 19 genes that are implicated in monogenic PD [3], 90 genetic risk variants have been found to contribute to disease susceptibility [5]. Carriers of a number of heterozygous mutations in the glucocerebrosidase gene (*GBA1*) have been shown to have a 10-30% chance of developing PD, making *GBA1* mutations the most common genetic risk factor for the disease [6, 7].

*GBA1* encodes for glucocerebrosidase (GCase), a lysosomal hydrolase that degrades glucosylceramide (GlcCer). The molecular weight of GCase ranges from between approximately 55-69 kDa, depending on the degree of glycosylation [8, 9]. In *GBA1-*associated PD, both loss-of-function and gain-of-function effects of *GBA1* mutations on GCase biology are involved in the cellular pathobiology (Reviewed by: [10]). Endoplasmic reticulum (ER) retention of misfolded GCase induces ER stress (a gain-of-function effect) and exacerbates the lysosomal GCase activity deficit (a loss-of-function effect) [11–14]. An increase in cellular α-synuclein increases ER stress and fragmentation [15] and further impairs ER-lysosomal GCase trafficking in a toxic cycle [15–17].

The relevance of *GBA1* to PD was first demonstrated in a series of clinical reports of Parkinsonism in patients with Gaucher’s disease (GD) [18–22], a lysosomal storage disorder caused by biallelic (either homozygous or compound heterozygous) *GBA1* mutations. This association was later confirmed in an international multicentre screening study [23]. Of the approximate 300 GD-causing *GBA1* mutations [24], 106 are associated with PD [25] and the penetrance of *GBA1* mutations in PD varies according to the type of mutation. Traditionally, *GBA1* mutations are classed as “risk” (e.g. E326K), “mild” (e.g. N370S) or “severe” (e.g. L444P) depending on their pathogenicity for GD in a homozygous state. Risk mutations do not cause GD, mild mutations cause type 1 (non-neuronopathic) GD and severe mutations cause type 2 or 3 (neuronopathic) GD [25]. Severe *GBA1* mutations confer a greater risk of developing PD and are associated with a younger age of onset, increased incidence of dementia, cognitive dysfunction, psychiatric symptoms and hyposmia [26–31]. The L444P mutation (c. 1448 T>C, exon 10) is the most common severe *GBA1* mutation in PD [23, 32, 33], although its frequency varies across populations [34, 35]. However, why the L444P mutation predisposes to more severe phenotypes remains unknown, and there are still many uncertainties surrounding mechanisms of GCase dysfunction in PD.

Transgenic rodent models have considerably enhanced our understanding of PD genetics by enabling biochemical and *in vivo* behavioural readouts of disease progression due to the addition of the human mutant gene [36]. The importance of studying human rather than mouse GCase dysfunction *in vivo* has been highlighted by studies demonstrating that mouse hepatic GCase activity is mouse strain-dependent [37], and that the efficacy of GCase activators varies between mouse and human GCase [38]. However, up to this point, no humanised models of *GBA*-associated PD were available and existing models have various limitations [39].

To bridge this gap, we have generated and characterised humanised transgenic mice that are hemizygous for either the human wildtype (hWT) or the human L444P mutant *GBA1* transgene on a mouse heterozygous *Gba^+/-^* background, named *Gba*^+/-^*GBA*^WT^ or *Gba*^+/-^*GBA*^L444P^, respectively. Using a BAC transgenic approach, site-specific integration of the hWT *GBA1* genomic locus including its endogenous promotor and flanking regulatory elements was achieved to generate hWT mice. Recombineering was used to generate a BAC genomic DNA *GBA1* transgene carrying the L444P point mutation which was then inserted into the same site in the genome to generate L444P mice. The use of the whole genomic locus inserted in a site-specific manner ensures consistent and correct spatiotemporal expression of the transgene across the hWT and L444P mice, making these genotypes directly comparable. Furthermore, our breeding pairs produce *Gba* heterozygous mice (*Gba*^+/-^) [40] and wild-type mice as littermate controls of the humanised transgenic mice, which allows us to directly address the potential loss-of-function or gain-of-function effects of the L444P mutation in driving pathology. This novel model displayed several pathological phenotypes, including reduced GCase activity, ER-retention of mutant GCase, impaired striatal dopamine release and accumulation of α-synuclein oligomers.

## Methods

### Generating and genotyping *GBA1* transgenic mice

Humanised *GBA1* transgenic mouse lines were created by transfection of mouse embryonic stem (ES) cells with a BAC vector carrying a 24 kb genomic DNA insert containing either the full length *GBA1* hWT or the *GBA1* L444P mutation insert retrofitted with the PhiC31 integrase exchange machinery to enable site specific integration into the mouse *Gt(ROSA)26Sor* locus in (C57BL/6J x 129S6/SvEvTac)F1-derived IDG26.10-3 mouse embryonic stem cells as previously described [41]. To generate experimental mouse cohorts, mice hemizygous for either the wildtype or L444P human *GBA1* transgene were crossed with C57BL/6 mice heterozygous for mouse *Gba* [40] (Strain name: B6.129S6-Gba-tm1Nsb/J, Jackson Laboratories stock number: 003321) resulting in mice carrying a single copy of either the hWT or L444P gene and a single copy of the mouse *Gba* gene.

DNA was extracted from mouse ear-clip biopsies in lysis buffer (50 mM KCl, 1.5 mM MgCl_2_, 10 mM Tris pH 8.5, 0.45% IGEPAL CA-630, 0.45% Tween 20) supplemented with 0.56 mg/mL proteinase K. Samples were vortexed, incubated at 65°C for one hour and 95°C for a further 30 min and centrifuged at 2400 rcf for 1 min. DNA segments of interest were amplified by polymerase chain reaction (PCR) in the presence of relevant primer pairs (**Table 1**). PCR products were either (i) run on 1% agarose gels (**Table 1, Reaction A and C**), or (ii) incubated with the NciI restriction enzyme (0.25%) for one hour at 37°C, and subsequently run on a 2% agarose gel (**Table 1, Reaction B)**. This enzymatic digest reaction enables the distinction between the wildtype and L444P *GBA1* human transgene as NciI only cleaves the amplified DNA segment if the L444P mutation is present (recognition site: CC^SGG). Representative genotyping results are shown in **Suppl. Fig. 1 B-D**.

**Table 1:**
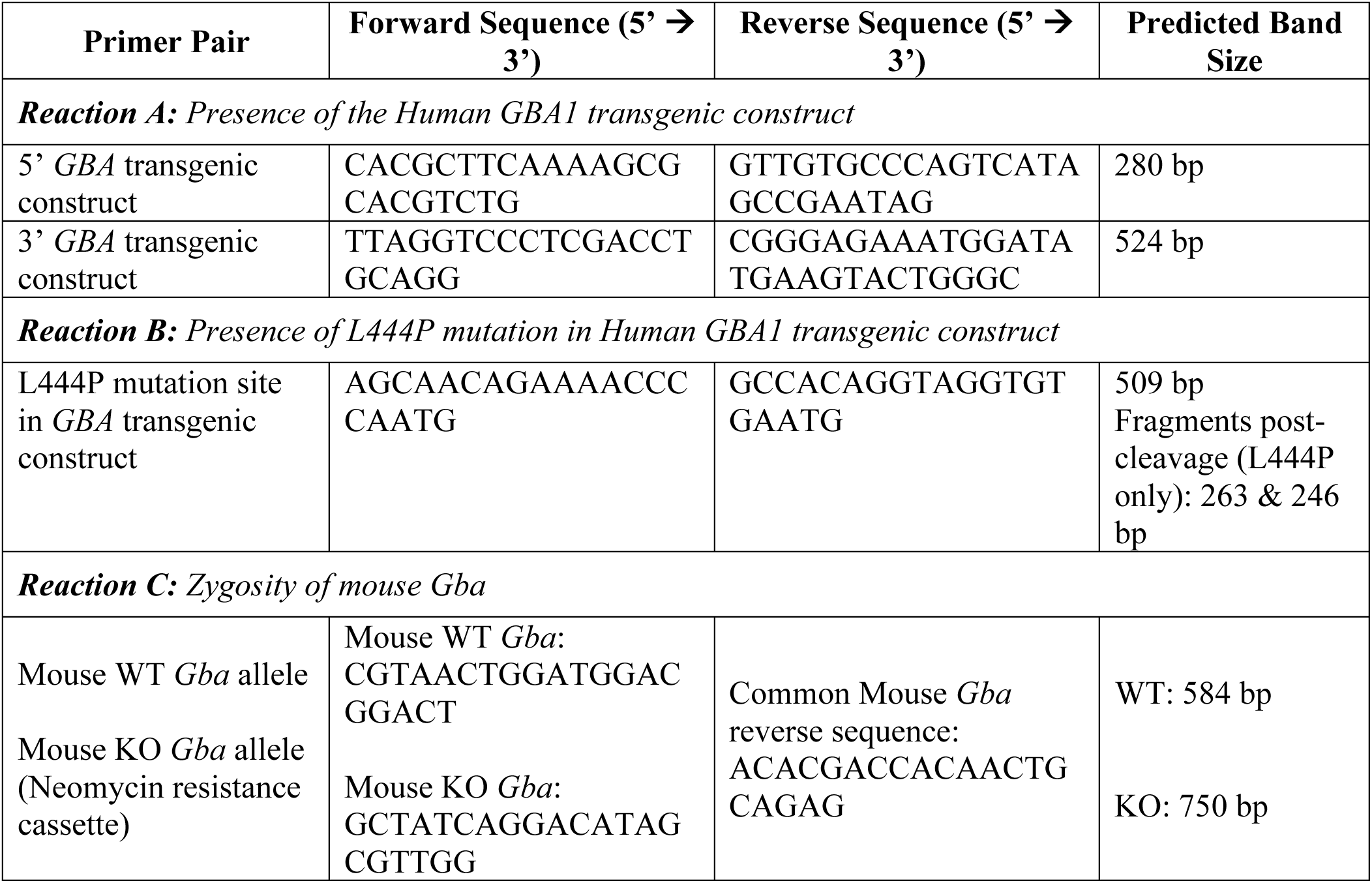
Primer sequences for genotyping *GBA* transgenic mice.

### Animal husbandry

Mice were maintained in accordance with UK Home Office regulations, under the Animals (Scientific Procedures) Act of 1986. They were housed at 20-24°C, 45-65% humidity on a 12:12 light cycle with 30 min dawn and 30 min dusk. Food and water were available *ad libitum*. Both male and female mice were used for all experiments.

### 4-MUG GCase activity assay

Dissected brain tissue was homogenised and sonicated on ice in citrate phosphate-based buffer (111 mM Na_2_HPO_4_, 44 mM C₆H₈O₇, pH 5.4) with 0.25% Triton X-100, 0.25% taurocholic acid and protease and phosphatase inhibitors (1 tablet per 10 mL solution). Samples were centrifuged at 1200 rcf for 5 min at 4°C and protein concentration of the supernatant was determined using a bicinchoninic acid (BCA) assay. Each sample (50 μL per well) was incubated with 20 μL of 10 mM 4-methylumbelliferyl β-D-glucopyranosidase (4-MUG) fluorescent substrate for 1 hour at 37°C. The reaction was quenched by adding 70 μL of 0.2 M glycine (pH 10.8) before fluorescence was read on a PHERAstar FSX microplate reader (BMG Labtech) at excitation/emission wavelengths of 350/450 nm. Negative control samples were pre-treated with 100 μM (final concentration) of conduritol B epoxide (CBE) (30 min at 37°C) prior to the addition of 4-MUG.

### Western blotting

Dissected brain tissue was homogenised in radioimmunoprecipitation assay (RIPA) buffer (50 mM Tris pH 8, 150 mM NaCl, 1% IGEPAL, 0.5% sodium deoxycholate, 0.1% SDS, supplemented with protease and phosphatase inhibitors at 1 tablet per 10 mL solution), centrifuged at 6000 rcf for 5 min at 4°C and protein concentration of the supernatant determined using a BCA assay. Samples were diluted as required in RIPA buffer and Laemmli sample buffer and denatured at 95°C for 5 min. Pre-cast polyacrylamide 4-15% gels were run and proteins transferred to PVDF membranes using the Trans-Blot Turbo Transfer System (Bio-Rad). Membranes were washed in 0.1% TBS-T, blocked in 4% milk for one hour at room temperature and subsequently treated with primary antibodies (GBA ab128879 Abcam; TH AB152 Millipore; SNAP-25 610366 BD Biosciences; VAMP2 v1389 Sigma; Synaptotagmin ab13259 Abcam; α-synuclein ab1903 Abcam; BiP 9956 Cell Signalling; LIMP2 4621 ProSi; LC3 2775 Cell Signalling; LAMP1 ab24170 Abcam; β-actin ab49900 Abcam) diluted in 4 % milk overnight at 4°C. Membranes were washed in 0.1% TBS-T, incubated with secondary antibodies at room temperature for one hour and washed in 0.1% TBS-T before imaging. β-actin was used as a loading control and was probed for with HRP anti- β-actin antibody, ab49900 Abcam (1:50,000) for one hour at room temperature. Membranes were developed with Immobilon western chemiluminescent HRP substrate (Millipore) and imaged with the Chemidoc MP Imaging System (Bio-Rad).

### Glycosphingolipid analysis

The midbrain and striatum were collected from a sacrificed mouse. Based on the mass of tissue, 25 mM Kpi-buffer pH 6.5 was added to obtain the following concentrations: 8 mg tissue/mL for midbrain and 1.5 mg/mL for striatum. Organs were lysed using 1.0 mm glass beads in an MP Biomedicals FastPrep-24. Protein levels were subsequently measured by BCA assay, then aliquoted and stored at −80°C until use.

Extractions and LC-MS/MS measurements of glycosphingoid bases and neutral glycosphingolipids were performed according to the method described previously [42]. Before extraction, 20 μL of ^13^C_5_-GlcSph (0.1 pmol/μL in methanol) and 20 μL of C17-dihydroceramide (1 pmol/μL in methanol) were spiked into 20 μL of midbrain lysates and into 10 μL of striatum lysates. Methanol and chloroform (2:1, v/v) were then added, and the samples were incubated for 30 min in the Eppendorf ThermoMixer at room temperature and 1200 rpm mixing frequency. The samples were centrifuged at 15,700 g for 10 min to pellet precipitated proteins, after which the supernatant was carefully transferred to a fresh tube. Chloroform and 100 mM ammonium formate buffer (pH 3.1) were added to the supernatant to obtain a final solvent ratio of the Bligh and Dyer procedure [43] 1:1:0.9 (methanol:chloroform: formate buffer), promoting two-phase separation. The upper phase containing lyso-glycosphingolipids was taken and concentrated at 45°C in an Eppendorf Concentrator Plus. The lower phase containing neutral glycosphingolipids was transferred to Pyrex tubes and evaporated to dryness at 45 °C with a gentle nitrogen stream. Then, deacylation was performed by adding 500 μL of 0.1 M NaOH in methanol using microwave-assisted saponification [44, 45]. After cooling, the samples were neutralised with 50 μL of 1 M HCl in methanol and dried again. The dried residues of both lyso-glycosphingolipids and neutral glycosphingolipids underwent butanol/water (1:1, v/v) extraction. Then, the water phase was discarded, and the butanol phase was dried and reconstructed in 100 μL of acetonitrile/methanol (9:1, v/v), mixed thoroughly, briefly sonicated in a water bath for 30 s, and centrifuged. The resulting supernatant was transferred to an autosampler vial for LC–MS/MS analysis. At last, 10 μL of the sample was injected into the UPLC–MS system.

Measurements were performed on a Waters UPLC–Xevo TQS micro system (Waters Corporation, Milford, MA) in positive-ion mode, using an electrospray ionisation (ESI) source. To separate glycosphingolipids containing glucosyl or galactosyl groups, analyses were performed using the previously described hydrophilic interaction liquid chromatography (HILIC) method [42]. Data were analysed with Masslynx 4.2 software (Waters Corporation). The compounds ^13^C_5_-GlcSph and C17-dihydroceramide were synthesised at the Department of Bio-organic Synthesis, Leiden Institute of Chemistry, Leiden University [46, 47]. Standards including GlcSph and GalSph, GlcCer (d18:1/16:0) and galactosylceramide (GalCer) (d18:1/16:0) were obtained from Avanti Polar Lipids (Alabaster, AL).

### Immunofluorescence

Mice were euthanized via intraperitoneal administration of a terminal dose of pentobarbitone. Following the loss of its tail-pinch, pedal and blink reflexes, the mouse underwent transcardiac perfusion of 1X PBS, followed by filtered 4% paraformaldehyde (PFA) solution (pH 7.4). Brains were extracted and post-fixed in 4% PFA for 24 hours at 4°C before being transferred to PBS with Azide for longer-term storage. Following fixation, brains were dehydrated by 10 min consecutive incubations in 95% ethanol, two 100% ethanol incubations, two histoclear incubations and three paraplast incubations before being embedded in paraffin. Sections were cut to 8 µm thick for immunofluorescence. For staining, samples were re-hydrated and heat-mediated antigen retrieval performed in 1X citrate buffer pH 6.0 (Abcam). Slides were blocked in 10% Donkey serum (diluted in 0.4% PBS-Tween (PBS-T)) for one hour at room temperature and subsequently incubated with primary antibodies overnight at 4°C (Sheep Tyrosine Hydroxylase, AB1542 Merck Millipore (1:250). After three washes in 0.4% PBS-T, samples were treated with secondary antibodies (Donkey anti-Sheep Alexa Fluor 488, A-11015 Invitrogen (1:1000);) for one hour at room temperature. Samples were re-washed three times in 0.4% PBS-T and mounted with FluorSave (Millipore). Sections were imaged using the FV1000 Confocal microscope (Olympus). Image analysis was performed in ImageJ.

For 3,3’-diaminobenzidine (DAB, Sigma) staining samples were rehydrated and antigen retrieval performed as above before being incubated in 10% H_2_O_2_ in PBS, to reduce background. Samples were blocked in 10% serum in PBS-T and then incubated in primary antibody (TH ab152 Millipore 1:1000) at room temperature for one hour. Samples were washed and incubated with the biotinylated secondary antibody for one hour followed by further washes and application of ABC solution (Vector Laboratories) for 30 minutes. Samples were washed with PBS-T and DAB was applied to the samples in order to visualise the antibody. Samples were washed with dH_2_O twice and then serially dehydrated and taken to xylene before mounting with DPX.

### Proximity ligation assay (PLA)

PLA was performed as previously described in [48]. Briefly, experiments were performed using the Duolink kits (Sigma) according to manufacturer’s instruction. The mouse monoclonal anti-α-syn4D6, ab1903, Abcam, 1:2000), was used to prepare conjugates using Duolink Probemaker Plus and Minus kits. Following sample de-waxing and rehydration antigen retrieval was performed by microwaving samples in citrate buffer pH 6.0 (Abcam). Samples were blocked in 10% serum in TBS-T (0.05% Tween) for one hour followed by a one-hour room temperature incubation with primary antibody (TH ab76442 Abcam). Samples were then washed in TBS-T before incubation for one hour at room temperature with the secondary antibody (Alexa488, Life Technologies) after which they were washed with TBS-T. To perform PLA samples were then blocked in Duolink block solution at 37°C for one hour followed by incubation with α-syn conjugates diluted in Duolink PLA diluent overnight at 4°C. Samples were washed using TBS-T and ligation reagents for 1 hour at 37°C followed by four TBS-T washes and a 2.5-hour incubation at 37°C with Duolink amplification reagents. Samples were then washed and mounted using FluorSave (Calbiochem). PLA puncta were counted in 35 TH-positive cells of the SNpc per animal. For cortical quantification images were taken in the motor cortex and the PLA spot count was automated using ImageJ. For each animal counts were averaged over four cortical images. For all image analysis the animal ID and genotype were blinded.

## Stereology

TH-positive neurons of the SNpc were counted as previously described [49–51]. Briefly, the borders of the SNpc and ventral tegmental area (VTA) were outlined using a distribution atlas of TH-positive cells. The number of SNpc TH-positive cells was counted on every fifth section of the serially sectioned mouse brain. Every TH-positive cell with a clear nucleus was counted through the whole region. A mean was then calculated for each animal and used with the Abercrombie correction factor [52] to obtain an actual number of TH-positive cells in the structure. All imaging and counts were performed with the researcher blinded to genotype.

## Stereotactic α-synuclein PFF injections

Mice were aged to 3 months and received bilateral 1.5 μL mouse α-synuclein preformed fibrils (PFFs) by stereotactic injection into the striatum. Coordinates of +0.5 mm anteroposterior, +/- 2.5 mm mediolateral and -2.7 mm dorsoventral were used and PFFs were delivered at a flow rate of 200 nL/min using a 32 gauge Hamilton syringe (Hamilton) which was withdrawn 5 minutes post injection. Mice were aged for a further 3 months before being transcardially perfused with 4% PFA. The brain was removed and postfixed overnight in 4% PFA before being dehydrated and paraffin embedded as described above. Mouse PFFs were produced as previously described [48]. The entire SNpc was sectioned at 8 μm and stained with p-α-synuclein (EP1536Y, Abcam) and TH (sc-25269, Santa Cruz) and p-α-synuclein aggregates were counted in TH positive cells in every tenth section.

### Imaging

All images were taken using either the EVOSflAUTO (Thermo) or a laser scanning confocal microscope (FV1000, Olympus). Quantification was done manually (stereology, SNpc PLA) or analysed using ImageJ (striatal TH, cortical PLA) or Cell Profiler (SNpc LAMP1). In all cases, apart from immunofluorescence of striatal TH, the researcher was blinded to genotype and animal ID.

### Fast-scan cyclic voltammetry

Mice were euthanized by cervical dislocation and brains immediately extracted and immersed in ice-cold, HEPES-buffered solution (120 mM NaCl, 5 mM KCl, 20 mM NaHCO_3_, 6.7 mM HEPES acid, 3.3 mM HEPES sodium salt, 2 mM CaCl_2_, 2 mM MgSO_4_(7H_2_O), 1.25 mM KH_2_PO_4_, and 10 mM glucose). 300 μm coronal striatal sections were prepared using a vibratome (VT1200 S, Leica) and allowed to recover in room-temperature HEPES-buffered solution for at least 1 hour before transfer to the recording chamber. In the recording chamber, slices were maintained between 31-33°C and were superfused with bicarbonate-buffered artificial cerebrospinal fluid (aCSF) (26 mM NaHCO_3_, 1.3 mM MgSO_4_(7H_2_O), 124 mM NaCl, 3.8 mM KCl, 1.2 mM KH_2_PO_4_, 10 mM Glucose, 2.4 mM CaCl_2_(2H_2_O)) at a flow rate of 2.25 mL/min. All solutions were saturated with 95% O_2_/ 5% CO_2_. Slices were left to acclimatise in the recording chamber for a minimum of 20 min before recording.

The extracellular concentration of DA [DA]_o_ was measured using fast-scan cyclic voltammetry (FCV) in conjunction with a single-use carbon-fibre microelectrode (CFM) (tip length: 50-100 μm, diameter: 7 μm, fabricated in-house) that was connected to a Millar voltammeter (Julian Millar, Barts and the London School of Medicine and Dentistry). The voltammeter passed a biphasic triangular voltage waveform (-0.7 to +1.3 V vs an Ag/AgCl reference electrode) at a scan rate of 800 V/s and sampling frequency of 8 Hz across the CFM, as described previously [53, 54]. The background signal was subtracted by the voltammeter and the resultant current was digitized using a Digidata 1440A (Molecular Devices). To stimulate DA release, a concentric bipolar Pt/Ir electrode (diameter: 25 μm, FHC) was placed on the tissue surface approximately 100 μm from the CFM. This electrode delivered electrical pulses (amplitude: 0.65 mA, duration: 200 μs) at 2.5 min intervals. Evoked currents were confirmed as DA due to the potentials of their characteristic oxidation and reduction peaks at 500-600 mV and -200 mV, respectively. The oxidation peak of each voltammogram was measured from the baseline and plotted versus time to obtain readouts of [DA]_o_ over time. Following experimental recordings, CFMs were calibrated by washing on a 2 μM DA solution in aCSF, made immediately before use from stock solution of 2.5 mM DA in 0.1 M HC1O_4_ stored at 4°C.

Recordings were made in the dorsolateral quadrant of the dorsal striatum (dorsolateral striatum, DLS) or the nucleus accumbens (NAc) core. Multisite experiments were conducted blind to genotype and utilised striatal sections from age-matched and sex-matched pairs of mice. In these experiments, pre-determined recording sites in DLS and NAc were sampled in a random order, alternating between two slices from different genotypes. For drug wash-on experiments all recordings were made in the DLS. Drug solutions (5 μM cocaine (Sigma-Aldrich); 44 nM and 1 μM quinpirole hydrochloride (1061, Tocris Bioscience); 10 μM bicuculine (ab120107, Abcam) and 4 μM CGP-55845 hydrochloride (ab120337, Abcam) were prepared in aCSF immediately before application. All drugs were dissolved in distilled water or dimethyl sulfoxide (DMSO) to make stock aliquots at 1,000–10,000 × final concentrations and stored at −20 °C. For experiments involving wash-on of cocaine and GABA antagonists (bicuculine and CGP-55845), recordings were made in aCSF containing 1 μM dihydro-β-erythroidine hydrobromide (DHβE) (2349, Tocris Bioscience). DHβE is an antagonist at nicotinic acetylcholine receptors and is used to inhibit the influence of cholinergic signalling on DA dynamics, as previously described [53, 55–57].

Data were acquired using Axoscope, version 11 (Molecular Devices) and analysed with locally written Microsoft Excel macros. Data are presented per observation (multisite experiments) or per experimental slice (drug wash-on experiments).

### High-performance liquid chromatography

The concentration of DA and 3,4-dihydroxyphenylacetic acid (DOPAC) in striatal tissue was quantified using high-performance liquid chromatography (HPLC). Tissue punches were taken from the caudate putamen (CPu) and NAc of 300 μm thick coronal sections (diameter: 2 mm and 1.2 mm, respectively) and suspended in 0.1 M HClO_4_. For each mouse, one CPu and one NAc punch were taken per hemisphere, across four hemispheres. Punches were sonicated on ice at 25 kHz for 20 s and centrifuged at 14,000 rcf for 15 min at 4°C. An autosampler (AS2057 plus, JASCO) injected the supernatant into a liquid mobile phase (13% HPLC-grade MeOH, 0.12 M NaH_2_PO_4_, 0.8 mM EDTA, 0.5 mM OSA, pH 4.6) that was flowing at 1 mL/min. Analytes were separated across a column (Microsorb 100 - 5 C18, S150 x 4.6, Agilent) and were subsequently detected using a DECADE II SDC electrochemical detector (Antec Scientific), fitted with a glassy carbon working electrode set at +0.7 V with respect to an Ag/AgCl reference electrode. HPLC chromatograms were analysed with Clarity software (DataApex).

### Behavioural analysis

Behavioural testing was performed blind to genotype.

#### The CatWalk XT gait analysis system

Automated gait analysis was performed using the CatWalk XT gait analysis system (Noldus Information Technology). This system uses green LED lighting to reflect the mouse’s pawprints as it voluntarily transverses a 130 cm long glass walkway. The pawprints are recorded by a high speed camera in real-time and used to automatically generate a variety of gait parameters. In our setup, runs were classified as compliant if they lasted between 0.5-5 s and if the maximum variation in speed was less than 35%. Up to five compliant runs were acquired per mouse and the average data for each mouse was exported for analysis with the CatWalk XT software.

#### Spontaneous locomotor activity in the open field

Spontaneous locomotor activity (LMA) was measured using the Photobeam Activity System-Home Cage (PAS-HC) (San Diego Instruments). The apparatus consisted of a transparent acrylic container, surrounded by a 4 x 8 photobeam frame. Each container’s floor was covered with a thin layer of wood chips. Mice were placed in separate containers and their voluntary movements were tracked with photobeam breaks. Trials ran for 4 hours, from 9:00 to 13:00, and beam break data was collated into 15 min intervals. Mice did not have access to food or water during the trial and each mouse was tested once.

#### Accelerating rotarod

Mice’s coordination and balance were assessed using an ENV-577M rotarod (Med Associates). Mice were placed on the rotarod facing away from the experimenter and opposite to the direction of rotation. In each trial, the rotarod accelerated from 2.5 to 25 rpm over 5 min and the latency to fall was recorded. Mice underwent 3 trials per day over 5 consecutive days. Days 1-3 were considered as acclimatisation days and performance across trials on days 4-5 was averaged and used for genotype-comparison.

#### Vertical pole test

Coordination and speed were examined using the vertical pole test. A 60 cm long wooden pole was secured to the floor of a cage resembling the home cage. The pole was wrapped in bandage gauze to aid grip and the base of the pole was covered in wood chips. The mice were positioned within approximately 5 cm from the top of the pole with their head pointing toward the ceiling. The time that mice took to complete a t-turn (rotate their body to point downwards) and the time taken to walk down the pole were measured. The sum of these measures equalled the total descent time. Mice underwent 3 trials per day over 3 consecutive days. Days 1-2 were considered acclimatisation days and performance on day 3 was used for genotype-comparison. Trials were excluded from the analysis if the total descent time exceeded 20 s or if the mouse climbed over the top of the pole.

#### Inverted screen test

Grip strength was assessed using the inverted screen test. Each mouse was positioned standing upright in the centre of an 1849 cm^2^ wire-mesh square grid that was held above a cushioned surface. The screen was surrounded by a 4 cm thick wooden frame and individual squares of the grid (1.44 cm^2^) were made from 1 mm diameter wire. The screen was then inverted by the experimenter so that the mice were holding on to the underside. The latency to fall from the screen was recorded if the mouse fell before the maximum test time of 60 s. All mice were tested once.

#### T-maze

Spatial working memory was assessed by measuring spontaneous alternation with a T-maze, using a protocol adapted from [58]. The maze dimensions can also be found in [58]. Briefly, the mouse was placed in the start arm of the maze and was allowed to voluntarily enter either goal arm. Once the mouse’s four paws had entered a goal arm, it was trapped in its chosen goal arm for 30 s by lowering a sliding door. The mouse was subsequently returned to the start arm, with all doors re-opened, and was allowed to make a second choice of goal arm. A spontaneous alternation was recorded if the mouse entered opposite goal arms on its first and second choice. The outcome of the trial (alternate/no alternate) was recorded across 6 trials (2 trials per day over 3 consecutive days). This was used to calculate the % alternation across trials: (Number of alternations/ Number of completed trials) * 100. Trials were terminated and excluded from the analysis if the mouse failed to move into a goal arm within 120 s on either their first or second choice. Mice that completed less than 5 out of the total of 6 trials across 3 days were excluded from analysis (n = 7).

## Data analysis and presentation

Statistical analysis was performed with GraphPad Prism 10 (GraphPad software). Normality was assessed using the Shapiro-Wilk test to determine whether parametric or non-parametric approaches should be used to assess differences between experimental groups. For comparisons involving two groups only, parametric data was analysed using either a two-tailed unpaired t-test or a two-tailed paired t-test, whereas non-parametric data was analysed using a Wilcoxon matched-pairs signed rank test. For comparisons across three or more experimental groups, parametric data was analysed using a one-way analysis of variance (ANOVA), two-way ANOVA or a two-way repeated measures ANOVA, whilst non-parametric data was analysed with a Kruskal-Wallis test. A Tukey’s or Dunn’s post-hoc test were conducted if significant effects were identified in the preceding ANOVA or Kruskal-Wallis test, respectively. The statistical tests used for individual experiments are specified in the figure legend. Statistical significance was determined at p < 0.05 for all analyses. Data are presented as mean ± SEM and each datapoint represents one mouse, unless otherwise stated in the figure legend.

## Results

### BAC transgenic humanised *GBA*-L444P mice demonstrate retention of GCase in the endoplasmic reticulum as a gain-of-function phenotype related to *GBA*-associated PD

A bacterial artificial chromosome (BAC) vector carrying a 24 kb genomic DNA insert containing the complete human *GBA1* genomic DNA locus (∼10 kb) flanked by ∼7 kb upstream and ∼7 kb downstream genomic DNA sequence was engineered by homologous recombination in *E. coli* to generate the T to C base pair change yielding the L444P coding mutation. Humanised *GBA1* transgenic mouse lines were created by transfection of mouse embryonic stem (ES) cells with the full length *GBA1* hWT or *GBA1* L444P mutation BAC insert retrofitted with the PhiC31 integrase exchange machinery to enable site specific integration into the mouse *Gt(ROSA)26Sor* locus (**Suppl. Fig. 1 A-B**). This genomic DNA transgene approach combined with site-specific integration ensures that the integration site of the transgene is consistent and occurs at single copy, ensuring the transgene expression is equivalent across genotypes, making the hWT and L444P mice directly comparable.

To generate experimental mouse cohorts, mice hemizygous for either the hWT or L444P human *GBA1* transgenes were crossed with C57BL/6 mice heterozygous for mouse *Gba* [40] resulting in mice carrying a single copy of either the hWT or L444P gene and a single copy of the mouse *Gba* gene; ie: *Gba*^+/-^*GBA*^WT^ and *Gba*^+/-^*GBA*^L444P^ . The breeding strategy used in this study also produced wild-type non transgenic (nTG) mice and mice that were heterozygous for mouse *Gba* (*Gba*^+/-^) (**Fig. 1 A**). Having *Gba*^+/-^ and nTG mice as littermate controls for L444P mice allows the opportunity to understand whether particular pathological phenotypes can be attributed to loss- or gain-of-function effects of the L444P mutation (**Fig. 1 A**).

**Figure 1:**
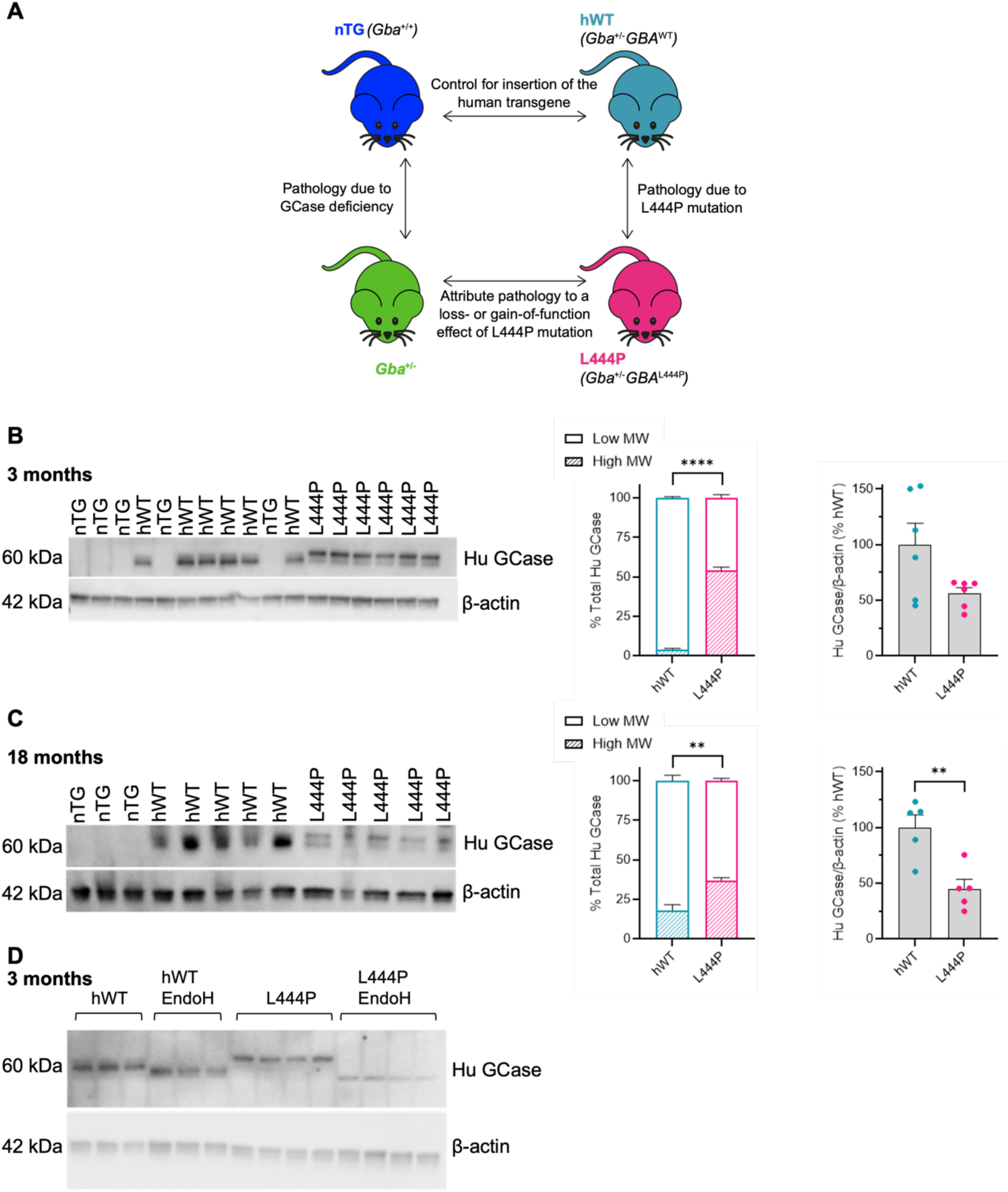
L444P mice express a high molecular weight, EndoH sensitive form of GCase, indicating entrapment of immature GCase in the ER. **(A)** Schematic of the four experimental genotypes used in this study and the questions that the genotype-comparisons can address. Western blots for human (Hu) GCase reveal an abundance of the higher molecular weight (MW) human GCase band in the midbrain of L444P mice aged 3 **(B)** and 18 **(C)** months old. The total levels of human GCase are significantly reduced in the L444P midbrain at 18 months. **(D)** Representative western blot illustrating a downward shift in the molecular weight of the human GCase band in L444P mice following EndoH treatment. **(B-C)** Two-tailed unpaired t-test: **(B)** 3 months high MW p < 0.0001, 3 months total p = 0.0527, **(C)** 18 months high MW p = 0.0013, 18 months total p = 0.0048. ** p < 0.01, **** p < 0.0001. n = 3-6 mice per group.

We first assessed human GCase expression levels across hWT and L444P mice by western blot. Total human GCase protein expression was reduced, although not significantly, between the hWT and L444P mice at 3 months-old. It was also apparent that there was a clear shift in the predominant molecular weight of the GCase protein expressed in the L444P mice compared to the hWT mice. The high molecular weight human GCase band was more abundant in L444P mice vs hWT mice, accounting for over 50% of total GCase present in 3 month-old mice (**Fig. 1 B**). In 18 month-old L444P mice, the high molecular weight human GCase band remained more abundant vs hWT mice; however, total human GCase levels were significantly lower in L444P than hWT mice at 18 months (**Fig. 1 C**).

Previous studies have indicated that various *GBA1* mutations can give rise to the accumulation of immature, misfolded GCase in the ER and thereby disturb GCase trafficking to the lysosome [11, 12, 59–61]. To assess whether the high molecular weight GCase band represented GCase that had not yet been processed through the Golgi, we treated tissue homogenates with EndoH to cleave high mannose oligosaccharides from the GCase protein [62]. Following EndoH treatment, we observed a downward shift in the molecular weight of the GCase band in mice carrying the L444P mutation (**Fig. 1 D**). As proteins that have passed through the Golgi apparatus are EndoH resistant [62], this result indicated that a substantial amount of human GCase in the L444P mouse midbrain had not yet been processed by the Golgi apparatus and was therefore retained in the ER, corroborating our previous findings in induced pluripotent stem cell (iPSC) derived DA neurons harbouring the *GBA*-N370S mutation [11]. It has previously been shown that GCase retention in the ER can induce ER stress [11, 12, 14, 15, 63]. However, midbrain levels of BiP were not significantly different across L444P vs hWT mice by western blot at 3 and 18 months (**Suppl. Fig. 2 A**).

### BAC transgenic humanised *GBA*-L444P mice demonstrate reduced GCase activity in the brain as a loss-of-function phenotype related to *GBA*-associated PD

Having demonstrated the L444P mutation causes GCase retention, we next sought to investigate whether our L444P mice display loss-of-function phenotypes; namely whether GCase activity in the midbrain is impaired. Using the 4-MUG GCase activity assay in midbrain homogenates from 3 and 18 month-old mice, we found a progressive reduction in GCase activity with age in the L444P mice (**Fig. 2 A**). At 3 months GCase activity decreased by 26% in L444P mice compared to nTG, albeit not significantly. However, by 18 months, GCase activity was significantly reduced by 62% compared to nTG mice. As expected, we also observed a significant reduction in midbrain GCase activity of *Gba*^+/-^ mice at 3 and 18 months (**Fig. 2 A**). The GCase activity deficit in L444P mice was accompanied by increased LIMP2 levels in the midbrain at 3 months but not with age (**Suppl. Fig. 2 B**). On the other hand, no changes to the overall protein levels of LC3-II or LAMP1 were observed in the L444P midbrain at either age (**Suppl. Fig. 2 C-D**).

**Figure 2:**
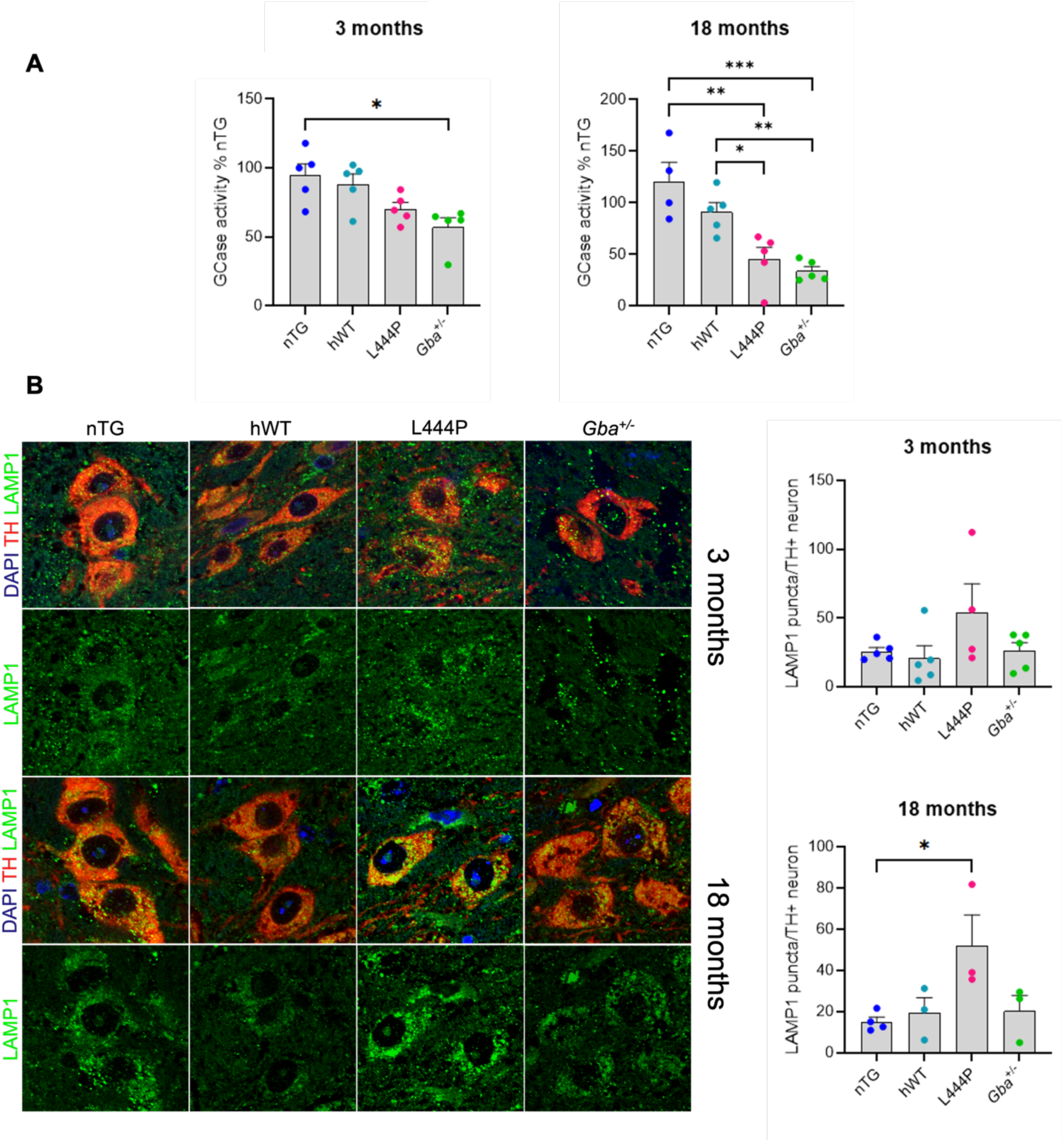
L444P mice display progressive GCase activity deficits and lysosomal dysfunction in the midbrain. **(A)** GCase activity is significantly impaired in the midbrain of L444P 18 month-old mice, measured by the 4-MUG GCase activity assay. (**B**) LAMP1 puncta are significantly increased in TH-positive neurons of the SNpc in L444P 18 month-old mice. Representative images of LAMP1 puncta in TH-positive neurons of the SNpc with quantification of the number of LAMP1 puncta shown in both 3 month-old and 18 month-old mice. (**A**) Kruskal-Wallis test with Dunn’s post-hoc test (3 months): Kruskal-Wallis statistic = 10.03, p = 0.0183, one-way ANOVA (Tukey’s) 3 month mice F = 5.983, one-way ANOVA with Tukey’s post-hoc test (18 months) F_(3,15)_ = 12.68, p = 0.0002. (**B**) One-way ANOVA (3 months): F_(3,15)_ = 1.886, p = 0.1754, one-way ANOVA with Tukey’s post-hoc test (18 months): F_(3,9)_ = 4.009, p = 0.0457. *p < 0.05, **p < 0.01, ***p < 0.001; n = 4-5 mice per genotype (**A**) and n = 3-5 mice per genotype (**B**).

Furthermore, the number of LAMP1 puncta per TH-positive neuron was modestly elevated in 3 month-old L444P mice and further became a significant increase in aged L444P mice (**Fig. 2 B**). This was only seen in the L444P mice and not in *Gba*^+/-^ mice suggesting this is a gain-of-function phenotype. No significant differences in the GCase glycosphingolipid substrate GlcCer were observed across genotypes in the midbrain or striatum at 3 months (**Suppl. Fig. 3 A**). On the other hand, midbrain and striatal glucosylsphingosine (GlcSph) levels were significantly increased in 3 month-old *Gba*^+/-^ mice vs nTG mice, although the one-way ANOVA did not detect an overall effect of genotype on GlcSph levels in these regions (**Suppl. Fig. 3 B**).

### SNpc DA neurons and striatal DA content are preserved in L444P mice

Having demonstrated that L444P mice display both gain-of-function and loss-of-function phenotypes associated with *GBA1* mutations in PD, we sought to assess whether this gives rise to DA neurodegeneration in the SNpc of aged L444P mice. We observed that stereological counts of tyrosine hydroxylase positive (TH+) neurons in the SNpc were not different across genotypes in 18 month-old mice (**Fig. 3 A**). This indicates that DA neurons are retained in L444P mice. Given that loss of striatal DA axons can precede midbrain DA neurodegeneration in PD [64], we subsequently investigated the abundance of DA axons in the striatum. We found that TH area in the CPu was similar across genotypes at 18 months of age, using immunofluorescence (**Fig. 3 B**). Moreover, 18 month-old L444P mice had comparable levels of striatal TH protein to other genotypes on a western blot (**Fig. 3 C**). In agreement with this, HPLC analysis showed that the tissue content of DA in the CPu (**Fig. 3 D**) or NAc (**Fig. 3 E**) was not different between genotypes in 3 month-old and aged (15-22 month-old) mice. Together, these results highlight that striatal DA axons are intact and DA levels are unchanged in L444P mice.

**Figure 3:**
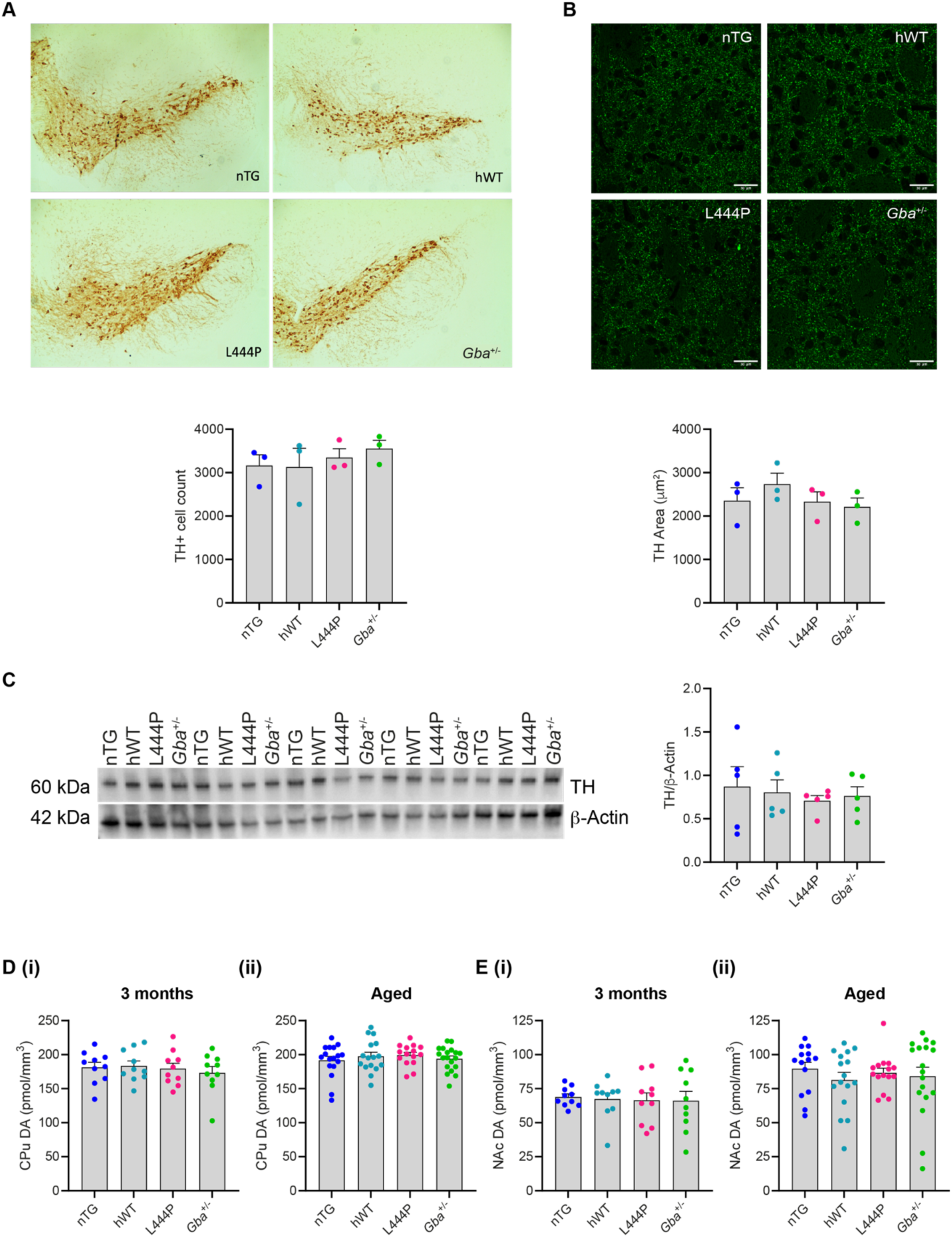
L444P mice do not display degeneration of SNpc dopaminergic neurons, striatal dopaminergic axons or loss of striatal dopamine content. **(A)** Stereological counts of TH-positive neurons in the SNpc were similar across genotypes at 18 months. (**B**) Representative immunofluorescent staining of striatal TH in 18 month-old mice and corresponding quantification showing no genotype-related differences in TH area. Scale bars = 30 μm. (**C**) Representative western blot and corresponding quantification showing comparable levels of TH across genotypes. (**D-E**) HPLC analysis using striatal tissue punches from 3 month-old and aged (15-22 month-old) mice showed that the DA content of the caudate putamen (CPu) (**D**) and nucleus accumbens (NAc) (**E**) is similar across genotypes. (**A-C, (D(i)**) One-way ANOVA: (**A**) F_(3,8)_ = 0.4655, p = 0.7143, (**B**) F_(3, 8)_ = 0.8352, p = 0.5113, (**C**) F_(3,16)_ = 0.2177, p = 0.8827, (**D(i)**) F_(3, 36)_ = 0.2938, p = 0.8296. (**D(ii), E**) Kruskal-Wallis test: (**D(ii)**) Kruskal-Wallis statistic = 1.089, p = 0.7796, (**E(i)**) Kruskal-Wallis statistic = 0.1361, p = 0.9872, (**E(ii)**) Kruskal-Wallis statistic = 1.720, p = 0.6326. n = 3 mice per genotype (**A-B**), n = 5 mice per genotype (**C**), n = 10 mice per genotype (**D(i), E(i)**), n = 15-19 mice per genotype (**D(ii)**), n = 14-18 mice per genotype (**E(ii)**).

### Evoked DA release is impaired in the DLS of L444P mice

A number of functional deficits in DA release and homeostasis have been reported to occur in PD before the onset of neurodegeneration [65]. Indeed, we have previously shown that other humanised transgenic rodent models of PD have impaired striatal DA release before, or in the absence of, neurodegeneration [65–68]. We tested whether L444P mice also exhibit dysregulation of striatal DA transmission despite the absence of DA neuron loss. Fast-scan cyclic voltammetry (FCV) was used to measure the concentration of extracellular DA [DA]_o_ evoked by single pulse electrical stimulations with sub-second resolution. Signals were confirmed as DA due to the characteristic potentials of the DA oxidation and reduction peaks (-200 mV and 500-600 mV, respectively), as shown on the representative voltammogram (**Fig. 4A, top**). [DA]_o_ evoked by single-pulse electrical stimulations were sampled from three recording sites in the dorsolateral quadrant of the dorsal striatum (dorsolateral striatum, DLS), which is predominantly innervated by DA neurons from SNpc, and two recording sites in the nucleus accumbens (NAc) core, which is predominantly innervated by DA neurons from the ventral tegmental area (VTA) (**Fig. 4 A, bottom**). Acute striatal brain sections from pairs of age- and sex-matched mice of different genotypes were simultaneously held in the recording chamber. Sampling was alternated across sections at each recording site, which enabled [DA]_o_ to be directly compared across genotypes. Firstly, we found that DLS [DA]_o_ (**Suppl. Fig. 4 A-B**) and NAc [DA]_o_ (**Suppl. Fig. 4 C-D**) were comparable in nTG vs hWT mice at 3 months and with age (18-22 months). This confirmed that insertion of the human BAC transgenic construct does not alter [DA]_o_. Next, we also observed that peak [DA]_o_ in the DLS (**Fig. 4 B-C**) and NAc (**Fig. 4 D-E**) at 3 months and old age (18-22 months) were similar in nTG compared to *Gba*^+/-^ mice. This indicates that impaired GCase activity in the *Gba*^+/-^ mice (**Fig. 2 A**) does not result in attenuated striatal [DA]_o_. However, [DA]_o_ was significantly lower in the DLS of L444P mice at 3 months and at 18-22 months (**Fig. 4 F-G**) compared to hWT mice. This deficit exclusively occurred in the DLS: L444P mice had similar NAc [DA]_o_ to hWT mice at 3 months (**Fig. 4 H**), and intriguingly, had relatively higher NAc [DA]_o_ at 18-22 months (**Fig. 4 I**). By contrast, evoked [DA]_o_ in DLS in L444P at 3 months were not significantly lower than in *Gba*^+/-^ mice (p = 0.06) (**Suppl. Fig. 4 E**). There were also no significant differences in DLS [DA]_o_ between aged L444P and *Gba*^+/-^ mice (**Suppl. Fig. 4 F**) or in NAc [DA]_o_ of L444P vs *Gba*^+/-^ mice at both ages (**Suppl. Fig. 4 G-H**). As a loss of GCase activity was not sufficient to alter [DA]_o_ in the DLS of *Gba*^+/-^ mice (**Fig. 4 B-C**) it is likely that the deficit in evoked [DA]_o_ in DLS of L444P mice is due conversely to a gain-of-function of GCase.

**Figure 4:**
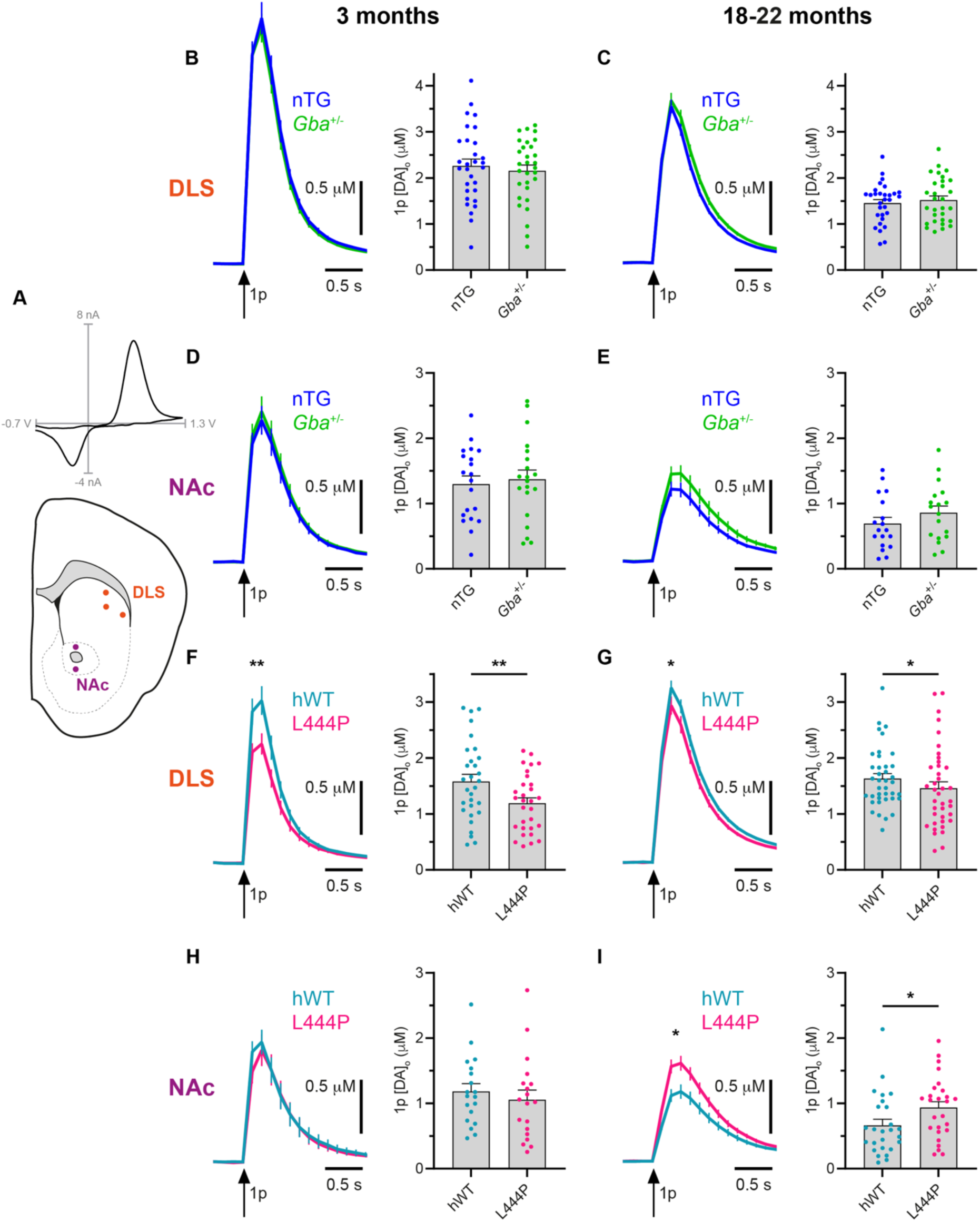
L444P mice, but not *Gba^+/-^* mice, have an evoked [DA]_o_ deficit exclusively in the DLS. (**A**) Representative DA voltammogram showing oxidation and reduction peaks at 0.5-0.6 V and -0.2 V, respectively, confirming that the signals are DA, and schematic of a coronal section illustrating the position of sampling sites in dorsolateral striatum (DLS) and nucleus accumbens core (NAc). (**B-I**) Mean [DA]_o_ transients evoked by single pulse electrical stimulations (↑ 1p) (left) and corresponding peak [DA]_o_ evoked by the same stimulations (right) in the DLS (**B, C, F, G**) and NAc (**D, E, H, I**) at 3 months (**B, D, F, H**) and 18-22 months (**C, E, G, I**). There were no differences in DLS [DA]_o_ of nTG and *Gba*^+/-^ mice at 3 months (**B**) or 18-22 months (**C**). (**D-E**) NAc [DA]_o_ was also not different across nTG and *Gba*^+/-^ mice at either age. L444P mice had impaired evoked [DA]_o_ in the DLS at 3 months (**F**) and 18-22 months (**G**). However, L444P mice had similar NAc [DA]_o_ to hWT mice at 3 months (**H**) and elevated NAc [DA]_o_ with age (**I**). (**B-F, H**) Two-tailed paired t-test: (**F**) p = 0.0012, (**G, I**) Wilcoxon matched- pairs signed rank test: (**G**) p = 0.0213, (**I**) p = 0.0102, all other graphs are p > 0.05. * p < 0.05, **p < 0.01. Each datapoint represents one recording site; (**B-C, F**) n = 30 recordings from 5 mice per genotype, (**D**) n = 20 recordings from 5 mice per genotype, (**E**) n = 18 recordings from 5 mice per genotype, (**G**) n = 39 recordings from 5 mice per genotype, (**H**) n = 19 recordings from 5 mice per genotype, (**I**) n = 26 recordings from 5 mice per genotype.

There are multiple mechanisms that could underpin this gain-of-function effect. We have already established that the [DA]_o_ deficit in L444P mice is not associated with SNpc DA neuron loss, or loss of striatal DA (**Fig. 3**). Dysregulation of many local striatal modulators of DA release have previously been reported in acute striatal slices and cultured DA neurons from other rodent models of PD [53, 69–71]. Therefore, we next asked if key regulators of DA dynamics are altered in L444P mice and if this could be associated with impaired [DA]_o_ in these mice. Using FCV, we measured the change in DLS [DA]_o_ in response to inhibition of the dopamine transporter (DAT) with cocaine (**Suppl. Fig. 5 A-B**), antagonism of GABA_A_ and GABA_B_ receptors (GABA_A+B_R) (**Suppl. Fig. 5 C-D**) or activation of dopamine D_2_ receptors (D_2_R) (**Suppl. Fig. 5 E-F**). We found that [DA]_o_ changed to a similar extent across genotypes following pharmacological modulation of these regulators in mice aged 3-6 months (**Suppl. Fig. 5 A, C, E**) and 15-22 months (**Suppl. Fig. 5 B, D, F**). Together, these results indicate that reduced evoked [DA]_o_ in DLS of L444P mice is not associated with changes to DAT function, tonic GABAergic inhibition of DA release or D_2_R-mediated inhibition of DA release. This may suggest that the dysfunctional changes mediating the [DA]_o_ deficit are due to other intrinsic features regulating DA releasability form DA axons. However, DA turnover appeared normal in L444P mice, as the DOPAC:DA ratio was similar across genotypes at 3 months and 15-22 months (**Suppl. Fig. 6 A**). Furthermore, western blots using striatal tissue showed no significant differences in the levels of key components of the synaptic release machinery: SNAP-25, VAMP2 and synaptotagmin across genotypes at 3 and 18 months (**Suppl. Fig. 6 B-D**).

### Reduced GCase activity is associated with α-synuclein oligomerisation in L444P mice

Given that α-synuclein accumulation and oligomerisation are key pathological features of *GBA*-associated PD and are associated with synaptic dysfunction, we next moved on to look at α-synuclein pathology. We first assessed total α-synuclein expression levels in both the midbrain and cortex of 3 month and aged mice but observed no changes across genotypes (**Fig. 5 A-B**).

**Figure 5:**
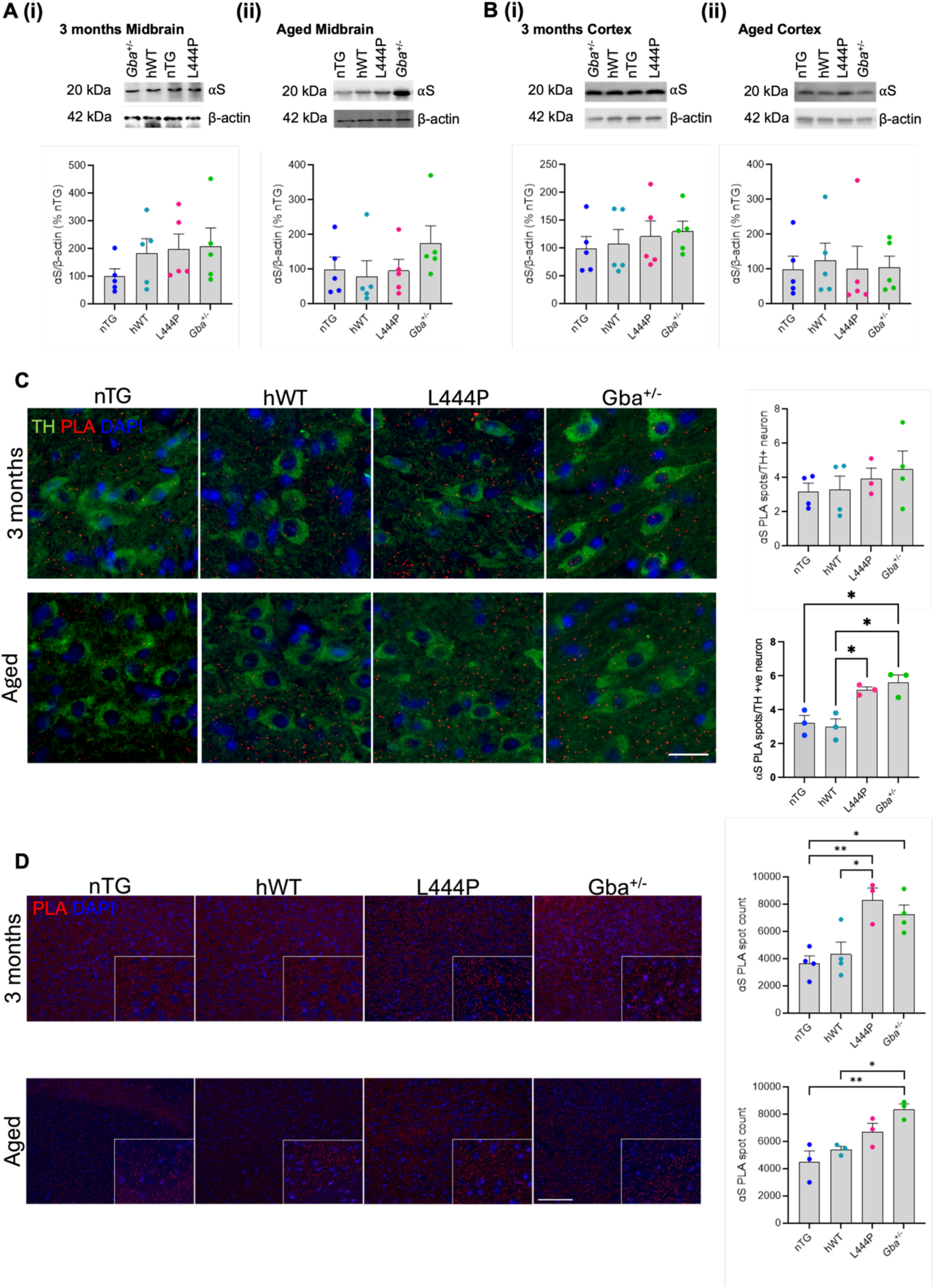
L444P mice accumulate α-synuclein oligomers in the midbrain and cortex. Total α-synuclein (αS) across genotypes on western blots of (**A**) midbrain and (**B**) cortex in 3 month and aged mice. Aged midbrain samples were taken from 18 month-old mice and aged cortex samples were from 15-21 month-old mice. αS is normalised to its own β-actin per mouse and expressed as a percentage of the mean αS levels in nTG mice. (**C**) Representative images of the Proximity Ligation Assay (PLA) signal in the TH positive cells of the SNpc. 35 neurons were quantified and averaged per mouse. (**D**) Representative images of the PLA in the motor cortex of 3 month and aged mice. (**A(i), C(3 months)**) One-way ANOVA: (**A(i)**) F_(3,16)_ = 0.9438, p = 0.4427, (**C(3 months)**) F_(3,11)_ = 0.6289, p = 0.6114, (**A(ii), B**) Kruskal-Wallis test: (**A(ii)**) Kruskal-Wallis statistic = 4.412, p = 0.2203, (**B(i-ii)**) Kruskal-Wallis statistic = 1.949, p = 0.5831, (**C**) Kruskal-Wallis with uncorrected Dunn’s test post-hoc test: H = 8.538, p = 0.0084 (aged)., (**D**) One-way ANOVA with Tukey’s post-hoc test: (3 months) F _(3,11)_ = 8.473, p = 0.0034, (aged) F _(3,8)_ = 9.222, p = 0.0056. *p < 0.05, **p < 0.01; (**A-B**) n = 5 mice per genotype, (**C-D**) n = 3-4 mice per genotype.

Given that the accumulation of oligomeric form of α-synuclein is considered an important part of early-stage pathology, we next assessed the levels of oligomeric α-synuclein in the DA neurons of the SNpc. There were no significant differences in oligomeric α-synuclein across genotypes in mice at 3 months of age; however, in aged mice we saw a significant increase in α-synuclein oligomers in both L444P and *Gba*^+/-^ mice compared to controls (**Fig. 5 C**). Given the known association of *GBA1* variants with an increased risk of cognitive symptoms and dementia in PD [26, 29, 72–76] we also assessed oligomeric α-synuclein in the cortex (**Fig. 5 D**). Here we found a significant increase in L444P and *Gba*^+/-^ mice at 3 months of age; however, in aged mice this increase was seen only in the *Gba*^+/-^ mice.

Taken together, this demonstrates that the accumulation of α-synuclein is associated with a loss-of-function role of mutant GCase. To understand if the *GBA*-L444P mutation was capable of driving α-synuclein induced cell loss or exacerbating α-synuclein pathology we performed striatal injection of α-synuclein preformed fibrils (PFFs) in mice. We observed no increase in the number of α-synuclein SNpc inclusions or DA neuron loss in either the L444P or *Gba*^+/-^mice **(Suppl. Fig. 7)** injected with α-synuclein PFFs.

### Behavioural phenotyping reveals aged L444P mice have mild gait deficits

So far we established that distinct pathological features in L444P mice can be attributed to gain-of-function effects of the mutation, such as protein glycosylation and striatal DA release deficits, and loss-of-function effects, such as reduced GCase function and α-synuclein oligomerisation. We next asked if these disease-relevant changes resulted in behavioural phenotypes in the absence of SNpc dopaminergic neurodegeneration. To address this, we assessed various parameters of spontaneous motor and non-motor function in a cohort of aged mice (15-21 months). Firstly, the CatWalk XT automated gait analysis system was used to record the gait of the mice as they transversed a glass walkway. Although the overall speed, forelimb swing duration and forelimb swing speed of the mice were similar across genotypes (**Fig. 6 A, Suppl. Fig. 8 A-B**), L444P mice had a significantly longer hindlimb swing duration and slower hindlimb swing speed (**Fig. 6 B-C**). These gait deficits were not accompanied by altered spontaneous locomotor activity (LMA) as the number of photobeam beam breaks made by the mice in the open field was similar across genotypes (**Fig. 6 D**). We also investigated coordination in the mice by measuring their latency to fall from an accelerating rotarod (**Fig. 6 E, Suppl. Fig. 8 C**) and time to descend a vertical pole (**Fig. 6 F**). No differences were observed in the rotarod performance averaged across test days (**Fig. 6 E**) or normalised to Day 1 (**Suppl. Fig. 8 C**). There were also no genotype-related differences in the total time to descend the vertical pole (**Fig. 6 F**), the time to make a t-turn (**Suppl. Fig. 8 D**) or the walk-down time (**Suppl. Fig. 8 E**). Moreover, no deficits in grip strength were observed across any of the genotypes using the inverted screen test (**Suppl. Fig. 8 F**); most of the mice held on to the inverted grid for the maximum trial time of 60 seconds. Finally, as PD patients with *GBA1* mutations are more likely to have cognitive deficits and dementia [26, 29, 72–76], we tested spatial working memory in the mice by assessing spontaneous alternations using the T-maze, as previously described [58]. However, the proportion of trials that resulted in an alternation was not significantly different across genotypes, indicating that L444P mice do not have impaired spatial working memory (**Suppl. Fig. 8 G**).

**Figure 6:**
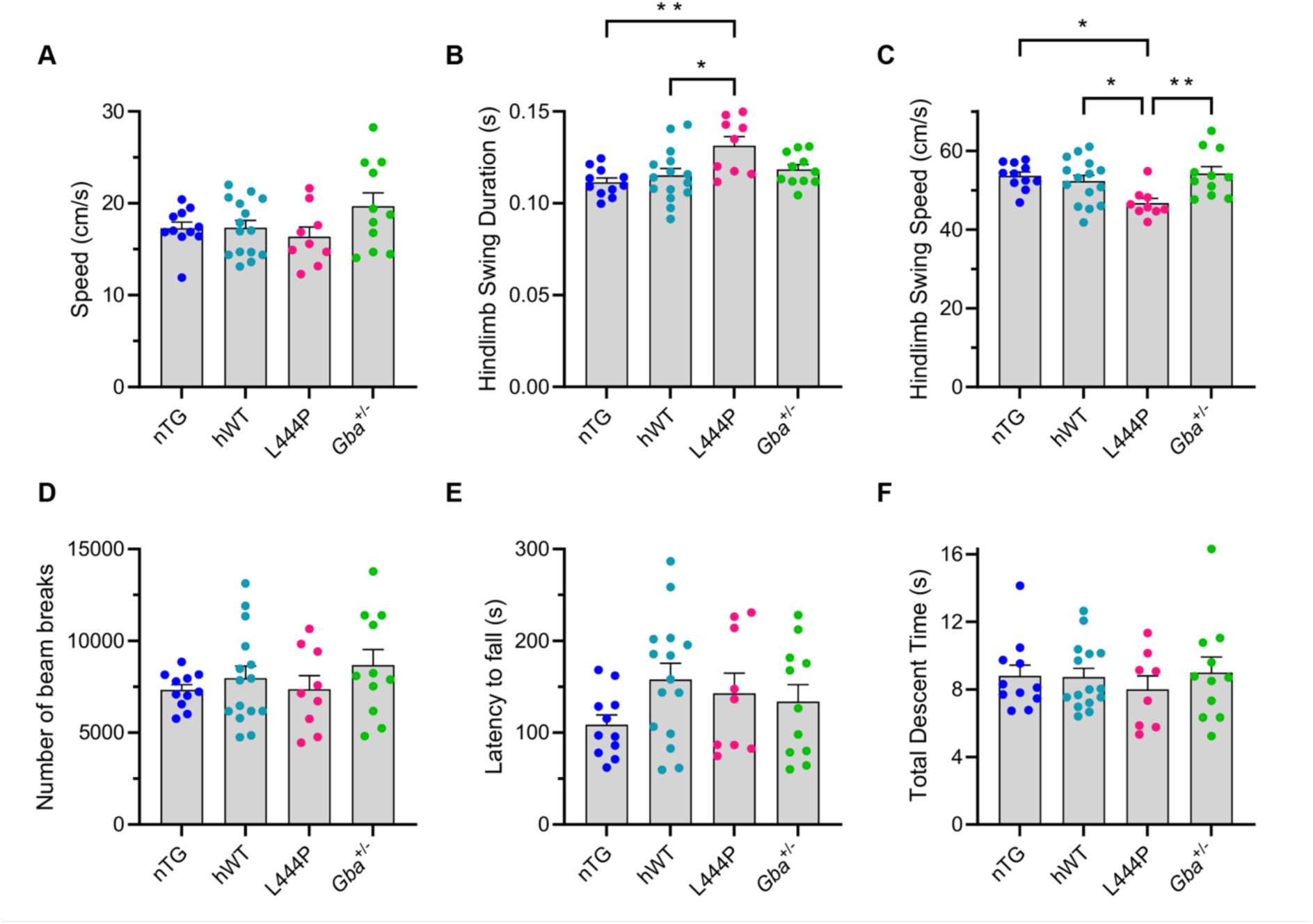
L444P mice have mild gait deficits but no changes to overall speed, locomotor activity and coordination. (**A-C**) Using the CatWalk XT automated gait analysis system, overall speed (**A**) was similar across genotypes; however, L444P mice displayed a significantly longer hindlimb swing duration (**B**) and significantly reduced hindlimb swing speed (**C**). (**D**) The number of photobeam breaks in the spontaneous locomotor activity test was comparable across genotypes. The latency to fall from the rotarod (**E**) and the total time taken to descend the vertical pole (**F**) were also not significantly different across genotypes. (**A**) one-way ANOVA: (**A**) F_(3, 42)_ = 1.814, p = 0.1592; (**B-C**) one-way ANOVA with Tukey’s post-hoc test: (**B**) F_(3, 42)_ = 5.056, p = 0.0044, (**C**) F_(3, 42)_ = 4.642, p = 0.0068, (**D-E**) one-way ANOVA: (**D**) F_(3, 42)_ = 0.7979, p = 0.5020, (**E**) F_(3, 42)_ = 1.489, p = 0.2315. (**F**) Kruskal-Wallis test: Kruskal-Wallis statistic = 0.7027, p = 0.8726. *p < 0.05 **p < 0.01; (**A-E**) n = 9-15 mice per genotype; (**F**) n = 8-15 mice per genotype.

## Discussion

It is widely recognized that *GBA1* mutations are the strongest genetic risk factor for PD but the mechanisms by which they increase PD susceptibility remain unclear [7]. The generation of genetic rodent models of *GBA1*-associated PD has proven challenging, and existing models have limited ability to recapitulate molecular and clinical features of PD [39]. We have generated and characterised the first humanised transgenic mouse model of *GBA1*-associated PD. BAC transgenesis allows the site-specific integration of the full length human *GBA1* gene, including all introns and exons, endogenous promotor and regulatory elements (**Suppl. Fig. 1 A**).

The heterozygous L444P mutation is the most prevalent severe *GBA1* mutation among PD patients [23, 32, 33]. This new *GBA-L444P* model offers advantages over currently available models of *GBA1*-associated PD. Modelling a PD-relevant *GBA1* mutation gives improved validity over pharmacological [77–79] models, genetic models that study homozygous *GBA1* mutations not generally seen in PD patients [77, 80–82], and models of GCase deficiency alone which lack gain-of-function phenotypes [40, 83, 84]. Previous models have introduced mutations to the endogenous mouse *Gba* gene [80, 81, 85–87]. However, hepatic GCase activity in mice is strain-dependent [37] and the efficacy of certain GCase activators can vary across human and mouse forms of the protein [38]. Studying the human GCase protein in a transgenic model circumvents these issues and has better predictive value for the development of pharmacological GCase modulators. Our study design, which includes *Gba*^+/-^ and wild-type nTG littermate controls, enables clarification on gain- and loss-of-function phenotypes.

We investigated GCase activity in *GBA-L444P* mice showing activity progressively declines with age in the midbrain, reaching a 62% reduction in mice aged 18 months (**Fig. 2 A**). Although not significant at 3 months, the magnitude of the loss of GCase activity was similar to previous observations in a heterozygous *Gba*-L444P mutant mouse [88–90]. Alongside this we observed a high abundance of Endo-H sensitive GCase in L444P mice (**Fig. 1 B-D**). Considering that glycosylated proteins that have been processed through the Golgi apparatus are Endo-H resistant, this phenotype indicates that immature, misfolded GCase is entrapped in the ER of the *GBA-L444P* mice [62]. This is the first demonstration of Endo-H sensitive GCase accumulation in a mouse model of *GBA1*-associated PD recapitulating similar phenotypes observed in human iPSC-derived DA neurons and midbrain organoid models, demonstrating this is an important feature in PD [11, 91]. Indeed, GCase chaperones, which show therapeutic promise for *GBA1*-associated PD, act to reduce ER-accumulation of mutant GCase by assisting its folding and lysosomal trafficking [92]. As this phenotype is evident in *GBA-L444P* mice at an early age (**Fig. 1 B**), it could be used as a biomarker of target engagement and compound effectiveness in preclinical drug development.

We observed a deficit in evoked dopamine release restricted to the DLS of *GBA-L444P* mice (**Fig. 4 F-G**), reflecting the PD-related preferential vulnerability and dysfunction of the nigrostriatal DA neurons that project from the SNpc to the DLS [93]. Considering that GCase deficiency did not lead to a significant deficit in evoked dopamine release in the DLS in the *Gba*^+/-^ mice (**Fig. 4 B-C**), it is apparent this is a gain-of-function effect of the L444P mutation. We established that this DA release deficit was not due to reduced DA turnover (**Suppl. Fig. 6 A**), SNpc DA neuron loss, attenuated striatal DA axon density or reduced striatal DA content (**Fig. 3**). We also confirmed there were no detected changes to DAT function, GABAergic inhibition of DA release or D_2_R-mediated inhibition of DA release (**Suppl. Fig. 5**). Therefore, it seems likely that this DA release deficit results from other changes to intrinsic mechanisms governing DA releasability. Although we did not find any gross changes in striatal levels of synaptic molecular machinery SNAP-25, synaptotagmin or VAMP2 (**Suppl. Fig. 6 B-D**), others have reported downregulation of genes implicated in synaptic vesicle recycling in cortex of *Gba*-*L444P/KO* mice [94]. Other studies have previously identified a reduction or re-distribution of synaptic vesicles as early disease phenotypes in various transgenic rodent models of PD [67, 94, 95].

Although we observed normal evoked dopamine release in *Gba*^+/-^ mice (**Fig. 4 B-E**), a previous study has reported reduced dopamine neurotransmission in mice treated with the GCase inhibitor CBE [77]. It should be noted that the techniques used to measure dopamine neurotransmission were not the same across studies and that CBE likely does not result in the same level of sustained GCase inhibition as genetic models, which may explain the variation in reported phenotypes. Interestingly, although evoked dopamine release in the DLS of L444P mice was lower than *Gba*^+/-^ mice, the reduction did not reach significance (**Suppl. Fig. 4 E-F**). We may therefore hypothesize that *Gba* haploinsufficiency could marginally reduce dopamine release in the DLS of *Gba*^+/-^ mice. This may serve as an explanation for why we did not observe significant differences in the *Gba*^+/-^ vs L444P mice.

We also observed a significant increase in dopamine release in the NAc of aged L444P mice (**Fig. 4 I**). Previously, increased striatal dopamine neurotransmission in α-synuclein overexpressing mouse models was attributed to compensatory effects to counter reduced release or impairments in autophagy or DAT function [96, 97]. Although we observed no significant changes to autophagy markers by western blot of midbrain tissue, we did see a significant increase in the number of LAMP1 puncta specifically in the dopaminergic neurons of the midbrain (**Suppl. Fig. 2, Fig. 2 B**). Rodríguez-Traver et al. (2024) [98] reported increased DA release in *GBA*-L444P iPSC-derived DA neurons, although the mechanistic basis of this phenotype was unclear. In any case, the opposing effects of the L444P mutation on dopamine release in the DLS and NAc in our model likely reflects differences between the populations of DA neurons (nigrostriatal and mesolimbic, respectively) that innervate these distinct striatal domains [99].

It is well established that α-synuclein accumulation in PD perturbs dopaminergic transmission before neurodegeneration and that GCase dysfunction and α-synuclein accumulation are thought to exacerbate each other in a mutually-toxic relationship [100, 101]. We previously developed the α-synuclein-PLA technique to detect early oligomeric α-synuclein species and have demonstrated its use in iPSC-derived DA neurons, α-synuclein overexpressing mice and post-mortem tissue from PD patients [48, 102, 103]. We identified a progressive accumulation of α-synuclein oligomers with age in the midbrain and the cortex of both L444P and *Gba*^+/-^mice (**Fig. 4 B-C**), highlighting that α-synuclein pathology is directly associated with loss of GCase activity. Interestingly, given that we did not observe a DA release deficit in *Gba*^+/-^ mice, who accumulated α-synuclein oligomers, it seems that α-synuclein accumulation is not the cause of the striatal DA release deficit in L444P mice. To the best of our knowledge, this is the first *in situ* demonstration of endogenous oligomeric α-synuclein accumulation in a mouse model of *GBA*-associated PD. Previously, the aggregation and spread of human α-synuclein oligomers were shown to be increased in *Gba*^+/L444P^ mice which virally overexpressed human α-synuclein, using a human α-synuclein-specific PLA [104].

Despite α-synuclein oligomer accumulation, total monomeric α-synuclein in various brain regions is unchanged in L444P mice (**Fig. 5 A-B**). This agrees with some [94, 105], but not all [90], studies investigating total α-synuclein in L444P mutant mice. In double mutant *hSNCA^A53T^Gba^+/L444P^* mice, α-synuclein was elevated in hippocampal neuronal cultures but not in brain tissue homogenates [105]. This likely reflects more effective degradation of α-synuclein via extra-neuronal mechanisms in the brain, which cannot be recapitulated in a neuronal culture system. This finding underscores the pathophysiological relevance of measuring α-synuclein directly from brain tissue of rodent models to examine the relationship between α-synuclein and GCase.

In our work nTG, L444P and *Gba*^+/-^ mice all displayed similar levels of p-α-synuclein inclusions and no dopaminergic cell loss in the SNpc following intrastriatal PFF injection (**Suppl. Fig. 7**), agreeing with some findings in other PD mouse models harbouring heterozygous *Gba* mutations. Previously, injection of α-synuclein PFFs did induce increased p-α-synuclein inclusions in the hippocampus of a heterozygous *Gba*-L444P mouse model of PD compared to wild-type mice, but not in the SNpc or cortex [88], while another study showed a significant increase of p-α-synuclein inclusions in the cortex of heterozygous *Gba*-L444P mice compared to wild-type mice after PFF injections [106]. In homozygous mouse models, injection of α-synuclein PFFs did not increase pathology in a *Gba1*-D409V knock-in model [107], but did in *Gba1*-E326K knock-in mice [85]. Our own α-synuclein-PLA data and PFF injection studies suggest that the loss of GCase may be important in driving the initial oligomerisation events of α-synuclein pathology, but not its later aggregation.

Consistent with other mouse models of *GBA*-associated PD [77, 80, 88, 90], we did not observe loss of SNpc DA neurons or loss of DA striatal axons in the L444P mice (**Fig. 3**). In PD patients, motor symptoms are not evident until at least 30% of SNpc neurons degenerate [108, 109]. Accordingly, behavioural deficits in the L444P mice were subtle; we observed a slower hindlimb swing speed but normal spontaneous locomotor activity and coordination (**Fig. 6**). On the other hand, *Gba*^+/-^ mice displayed no behavioural impairments indicating that gait deficits in the L444P mice do not directly arise due to loss of GCase activity. Previously it has been shown that mice harbouring L444P mutations, combined with induced α-synuclein pathology, spend less time rearing (indicative of hindlimb muscle weakness) [105] and have increased hindlimb clasping [94]. Considering that PD patients with *GBA1* mutations have a higher incidence of cognitive deficits and dementia [26, 29, 72–76], it was somewhat surprising that neither L444P nor *Gba*^+/-^ mice do not demonstrate impaired spatial working memory with the T-maze (**Suppl. Fig. 8 G**). However, our results corroborate previous reports of normal memory and normal spatial and associative learning in other models of *GBA*-associated PD [88, 110, 111].

In conclusion, we have generated and deeply-phenotyped a mouse that is heterozygous for the human *GBA*-L444P mutation, the first humanised transgenic model of *GBA1*-associated PD. Our breeding pairs produce *Gba*^+/-^ mice as littermate controls of L444P mice, enabling the investigation of loss- and gain-of-function effects of the *GBA-L444P* mutation. We show that *GBA-L444P* mice display early and sustained accumulation of immature GCase in the ER and a progressive loss of GCase activity. Ours is the first demonstration of impaired evoked dopamine release in the DLS in a mouse model of *GBA1*-associated PD through a gain-of-function mechanism of the *GBA-L444P* mutation, whereas the observed progressive accumulation of oligomeric α-synuclein occurs via a loss-of-function of GCase activity. This model represents a useful tool for future studies to elucidate *GBA*’s role in PD pathology, identify biomarkers of disease progression and develop disease-modifying therapies targeted at GCase function.

## Acknowledgements

The work was supported by the Monument Trust Discovery Award (J-1403) from Parkinson’s UK, Grant MJFF-023781 from the Michael J Fox Foundation, and Wellcome Collaborator Award 223202/Z/21/Z. TD was supported by a Wellcome Trust Neuroscience DPhil Studentship 102170/Z/13/Z.

## Author contributions

NCR, TD, JAA, BR, KB, NB-V, HW, MC, BD, KOB, MJCvdL and JMFGA designed and undertook experimental work, collecting and analysing data. JAA, NCR, SJC and RWM conceived the project. NCR, TD and RWM wrote, reviewed and edited the manuscript. RWM, SJC and NCR supervised the work.

## References

1. Obeso, J.A., et al., Past, present, and future of Parkinson’s disease: A special essay on the 200th Anniversary of the Shaking Palsy. Mov Disord, 2017. 32(9): p. 1264–1310.

2. Zhu, J., et al., Temporal trends in the prevalence of Parkinson’s disease from 1980 to 2023: a systematic review and meta-analysis. Lancet Healthy Longev, 2024. 5(7): p. e464–e479.

3. Deng, H., P. Wang, and J. Jankovic, The genetics of Parkinson disease. Ageing Res Rev, 2018. 42: p. 72–85.

4. Kalia, L.V. and A.E. Lang, Parkinson’s disease. Lancet, 2015. 386(9996): p. 896-912.

5. Nalls, M.A., et al., Identification of novel risk loci, causal insights, and heritable risk for Parkinson’s disease: a meta-analysis of genome-wide association studies. Lancet Neurol, 2019. 18(12): p. 1091–1102.

6. Anheim, M., et al., Penetrance of Parkinson disease in glucocerebrosidase gene mutation carriers. Neurology, 2012. 78(6): p. 417–20.

7. Balestrino, R., et al., Penetrance of Glucocerebrosidase (GBA) Mutations in Parkinson’s Disease: A Kin Cohort Study. Mov Disord, 2020. 35(11): p. 2111–2114.

8. Bergmann, J.E. and G.A. Grabowski, Posttranslational processing of human lysosomal acid beta-glucosidase: a continuum of defects in Gaucher disease type 1 and type 2 fibroblasts. Am J Hum Genet, 1989. 44(5): p. 741–50.

9. Erickson, A.H., E.I. Ginns, and J.A. Barranger, Biosynthesis of the lysosomal enzyme glucocerebrosidase. J Biol Chem, 1985. 260(26): p. 14319–24.

10. Yoon, J., C.Y. Lee, and V.S. Ah, Biochemical consequences of glucocerebrosidase 1 mutations in Parkinson’s disease. Neural Regen Res, 2024. 19(4): p. 725–727.

11. Fernandes, H.J., et al., ER Stress and Autophagic Perturbations Lead to Elevated Extracellular alpha-Synuclein in GBA-N370S Parkinson’s iPSC-Derived Dopamine Neurons. Stem Cell Reports, 2016. 6(3): p. 342–56.

12. Garcia-Sanz, P., et al., N370S-GBA1 mutation causes lysosomal cholesterol accumulation in Parkinson’s disease. Mov Disord, 2017. 32(10): p. 1409–1422.

13. Schondorf, D.C., et al., iPSC-derived neurons from GBA1-associated Parkinson’s disease patients show autophagic defects and impaired calcium homeostasis. Nat Commun, 2014. 5: p. 4028.

14. McNeill, A., et al., Ambroxol improves lysosomal biochemistry in glucocerebrosidase mutation-linked Parkinson disease cells. Brain, 2014. 137(Pt 5): p. 1481–95.

15. Stojkovska, I., et al., Rescue of alpha-synuclein aggregation in Parkinson’s patient neurons by synergistic enhancement of ER proteostasis and protein trafficking. Neuron, 2022. 110(3): p. 436–451 e11.

16. Mazzulli, J.R., et al., Gaucher disease glucocerebrosidase and alpha-synuclein form a bidirectional pathogenic loop in synucleinopathies. Cell, 2011. 146(1): p. 37–52.

17. Mazzulli, J.R., et al., *alpha-Synuclein-induced lysosomal dysfunction occurs through disruptions in protein trafficking in human midbrain synucleinopathy models*. Proc Natl Acad Sci U S A, 2016. 113(7): p. 1931–6.

18. Bembi, B., et al., Gaucher’s disease with Parkinson’s disease: clinical and pathological aspects. Neurology, 2003. 61(1): p. 99–101.

19. McKeran, R.O., et al., Neurological involvement in type 1 (adult) Gaucher’s disease. J Neurol Neurosurg Psychiatry, 1985. 48(2): p. 172–5.

20. Neudorfer, O., et al., Occurrence of Parkinson’s syndrome in type I Gaucher disease. QJM, 1996. 89(9): p. 691–4.

21. Tayebi, N., et al., Gaucher disease and parkinsonism: a phenotypic and genotypic characterization. Mol Genet Metab, 2001. 73(4): p. 313–21.

22. Varkonyi, J., et al., Gaucher disease associated with parkinsonism: four further case reports. Am J Med Genet A, 2003. 116A(4): p. 348-51.

23. Sidransky, E., et al., Multicenter analysis of glucocerebrosidase mutations in Parkinson’s disease. N Engl J Med, 2009. 361(17): p. 1651–61.

24. Hruska, K.S., et al., Gaucher disease: mutation and polymorphism spectrum in the glucocerebrosidase gene (GBA). Hum Mutat, 2008. 29(5): p. 567–83.

25. Parlar, S.C., et al., Classification of GBA1 Variants in Parkinson’s Disease: The GBA1-PD Browser. Mov Disord, 2023. 38(3): p. 489–495.

26. Cilia, R., et al., Survival and dementia in GBA-associated Parkinson’s disease: The mutation matters. Ann Neurol, 2016. 80(5): p. 662–673.

27. Gan-Or, Z., et al., Differential effects of severe vs mild GBA mutations on Parkinson disease. Neurology, 2015. 84(9): p. 880–7.

28. Lerche, S., et al., Parkinson’s Disease: Glucocerebrosidase 1 Mutation Severity Is Associated with CSF Alpha-Synuclein Profiles. Mov Disord, 2020. 35(3): p. 495–499.

29. Liu, G., et al., Specifically neuropathic Gaucher’s mutations accelerate cognitive decline in Parkinson’s. Ann Neurol, 2016. 80(5): p. 674–685.

30. Petrucci, S., et al., GBA-Related Parkinson’s Disease: Dissection of Genotype-Phenotype Correlates in a Large Italian Cohort. Mov Disord, 2020. 35(11): p. 2106–2111.

31. Thaler, A., et al., Parkinson’s disease phenotype is influenced by the severity of the mutations in the GBA gene. Parkinsonism Relat Disord, 2018. 55: p. 45–49.

32. Cook, L., et al., Parkinson’s disease variant detection and disclosure: PD GENEration, a North American study. Brain, 2024. 147(8): p. 2668–2679.

33. Westenberger, A., et al., Relevance of genetic testing in the gene-targeted trial era: the Rostock Parkinson’s disease study. Brain, 2024. 147(8): p. 2652–2667.

34. Gabbert, C., et al., GBA1 in Parkinson’s disease: variant detection and pathogenicity scoring matters. BMC Genomics, 2023. 24(1): p. 322.

35. Koros, C., et al., A Global Perspective of GBA1-Related Parkinson’s Disease: A Narrative Review. Genes (Basel), 2024. 15(12).

36. Vingill, S., N. Connor-Robson, and R. Wade-Martins, Are rodent models of Parkinson’s disease behaving as they should? Behav Brain Res, 2018. 352: p. 133–141.

37. Duran, A., et al., Identification of genetic modifiers of murine hepatic beta-glucocerebrosidase activity. Biochem Biophys Rep, 2021. 28: p. 101105.

38. Berger, Z., et al., Tool compounds robustly increase turnover of an artificial substrate by glucocerebrosidase in human brain lysates. PLoS One, 2015. 10(3): p. e0119141.

39. Farfel-Becker, T., et al., Can GBA1-Associated Parkinson Disease Be Modeled in the Mouse? Trends Neurosci, 2019. 42(9): p. 631–643.

40. Tybulewicz, V.L., et al., Animal model of Gaucher’s disease from targeted disruption of the mouse glucocerebrosidase gene. Nature, 1992. 357,(6377): p. 407-10.

41. Gordon, D., et al., Single-copy expression of an amyotrophic lateral sclerosis-linked TDP-43 mutation (M337V) in BAC transgenic mice leads to altered stress granule dynamics and progressive motor dysfunction. Neurobiol Dis, 2019. 121: p. 148–162.

42. Lelieveld, L.T., et al., Role of beta-glucosidase 2 in aberrant glycosphingolipid metabolism: model of glucocerebrosidase deficiency in zebrafish. J Lipid Res, 2019. 60(11): p. 1851–1867.

43. Bligh, E.G. and W.J. Dyer, A rapid method of total lipid extraction and purification. Can J Biochem Physiol, 1959. 37(8): p. 911–7.

44. Mirzaian, M., et al., Simultaneous quantitation of sphingoid bases by UPLC-ESI-MS/MS with identical (13)C-encoded internal standards. Clin Chim Acta, 2017. 466: p. 178–184.

45. Groener, J.E., et al., HPLC for simultaneous quantification of total ceramide, glucosylceramide, and ceramide trihexoside concentrations in plasma. Clin Chem, 2007. 53(4): p. 742–7.

46. Marques, A.R., et al., Glucosylated cholesterol in mammalian cells and tissues: formation and degradation by multiple cellular beta-glucosidases. J Lipid Res, 2016. 57(3): p. 451–63.

47. Wisse, P., et al., Synthesis of a Panel of Carbon-13-Labelled (Glyco)Sphingolipids. European Journal of Organic Chemistry, 2015. 2015(12): p. 2661–2677.

48. Bengoa-Vergniory, N., et al., CLR01 protects dopaminergic neurons in vitro and in mouse models of Parkinson’s disease. Nat Commun, 2020. 11(1): p. 4885.

49. Al-Wandi, A., et al., Absence of alpha-synuclein affects dopamine metabolism and synaptic markers in the striatum of aging mice. Neurobiol Aging, 2010. 31(5): p. 796–804.

50. Connor-Robson, N., et al., Combinational losses of synucleins reveal their differential requirements for compensating age-dependent alterations in motor behavior and dopamine metabolism. Neurobiol Aging, 2016. 46: p. 107–12.

51. Ninkina, N., et al., *beta-synuclein potentiates synaptic vesicle dopamine uptake and rescues dopaminergic neurons from MPTP-induced death in the absence of other synucleins*. J Biol Chem, 2021. 297(6): p. 101375.

52. Abercrombie, M., Estimation of nuclear population from microtome sections. Anat Rec, 1946. 94: p. 239–47.

53. Roberts, B.M., et al., GABA uptake transporters support dopamine release in dorsal striatum with maladaptive downregulation in a parkinsonism model. Nat Commun, 2020. 11(1): p. 4958.

54. Threlfell, S., et al., Striatal muscarinic receptors promote activity dependence of dopamine transmission via distinct receptor subtypes on cholinergic interneurons in ventral versus dorsal striatum. J Neurosci, 2010. 30(9): p. 3398–408.

55. Condon, M.D., et al., Plasticity in striatal dopamine release is governed by release-independent depression and the dopamine transporter. Nat Commun, 2019. 10(1): p. 4263.

56. Lopes, E.F., et al., Inhibition of Nigrostriatal Dopamine Release by Striatal GABA(A) and GABA(B) Receptors. J Neurosci, 2019. 39(6): p. 1058–1065.

57. Rice, M.E. and S.J. Cragg, Nicotine amplifies reward-related dopamine signals in striatum. Nat Neurosci, 2004. 7(6): p. 583–4.

58. Deacon, R.M. and J.N. Rawlins, T-maze alternation in the rodent. Nat Protoc, 2006. 1(1): p. 7–12.

59. Bendikov-Bar, I., et al., Characterization of the ERAD process of the L444P mutant glucocerebrosidase variant. Blood Cells Mol Dis, 2011. 46(1): p. 4–10.

60. Ron, I. and M. Horowitz, ER retention and degradation as the molecular basis underlying Gaucher disease heterogeneity. Hum Mol Genet, 2005. 14(16): p. 2387–98.

61. Schmitz, M., et al., Impaired trafficking of mutants of lysosomal glucocerebrosidase in Gaucher’s disease. Int J Biochem Cell Biol, 2005. 37(11): p. 2310–20.

62. Freeze, H.H. and C. Kranz, Endoglycosidase and glycoamidase release of N-linked glycans. Curr Protoc Mol Biol, 2010. **Chapter 17**: p. Unit 17 13A.

63. Gegg, M.E., et al., Glucocerebrosidase deficiency in substantia nigra of parkinson disease brains. Ann Neurol, 2012. 72(3): p. 455–63.

64. Burke, R.E. and K. O’Malley, Axon degeneration in Parkinson’s disease. Exp Neurol, 2013. 246: p. 72–83.

65. Cramb, K.M.L., et al., Impaired dopamine release in Parkinson’s disease. Brain, 2023. 146(8): p. 3117–3132.

66. Cramb, K.M.L., et al., *Dopamine neurotransmission in Parkinson’s disease*, in Handbook of Parkinson’s Disease Mechanisms, R. Moratalla and M.G. Murer, Editors. 2025, Elsevier. p. 171-189.

67. Janezic, S., et al., Deficits in dopaminergic transmission precede neuron loss and dysfunction in a new Parkinson model. Proc Natl Acad Sci U S A, 2013. 110(42): p. E4016–25.

68. Sloan, M., et al., LRRK2 BAC transgenic rats develop progressive, L-DOPA-responsive motor impairment, and deficits in dopamine circuit function. Hum Mol Genet, 2016. 25(5): p. 951–63.

69. Dagra, A., et al., *alpha-Synuclein-induced dysregulation of neuronal activity contributes to murine dopamine neuron vulnerability*. NPJ Parkinsons Dis, 2021. 7(1): p. 76.

70. Lin, M., et al., In Parkinson’s patient-derived dopamine neurons, the triplication of alpha-synuclein locus induces distinctive firing pattern by impeding D2 receptor autoinhibition. Acta Neuropathol Commun, 2021. 9(1): p. 107.

71. Threlfell, S., et al., Striatal Dopamine Transporter Function Is Facilitated by Converging Biology of alpha-Synuclein and Cholesterol. Front Cell Neurosci, 2021. 15: p. 658244.

72. Alcalay, R.N., et al., Cognitive performance of GBA mutation carriers with early-onset PD: the CORE-PD study. Neurology, 2012. 78(18): p. 1434–40.

73. Jesus, S., et al., GBA Variants Influence Motor and Non-Motor Features of Parkinson’s Disease. PLoS One, 2016. 11(12): p. e0167749.

74. Mata, I.F., et al., GBA Variants are associated with a distinct pattern of cognitive deficits in Parkinson’s disease. Mov Disord, 2016. 31(1): p. 95–102.

75. Straniero, L., et al., The SPID-GBA study: Sex distribution, Penetrance, Incidence, and Dementia in GBA-PD. Neurol Genet, 2020. 6(6): p. e523.

76. Szwedo, A.A., et al., GBA and APOE Impact Cognitive Decline in Parkinson’s Disease: A 10-Year Population-Based Study. Mov Disord, 2022. 37(5): p. 1016–1027.

77. Ginns, E.I., et al., Neuroinflammation and alpha-synuclein accumulation in response to glucocerebrosidase deficiency are accompanied by synaptic dysfunction. Mol Genet Metab, 2014. 111(2): p. 152–62.

78. Papadopoulos, V.E., et al., Modulation of beta-glucocerebrosidase increases alpha-synuclein secretion and exosome release in mouse models of Parkinson’s disease. Hum Mol Genet, 2018. 27(10): p. 1696–1710.

79. Mus, L., et al., Development and biochemical characterization of a mouse model of Parkinson’s disease bearing defective glucocerebrosidase activity. Neurobiol Dis, 2019. 124: p. 289–296.

80. Polinski, N.K., et al., Decreased glucocerebrosidase activity and substrate accumulation of glycosphingolipids in a novel GBA1 D409V knock-in mouse model. PLoS One, 2021. 16(6): p. e0252325.

81. Xu, Y.H., et al., Viable mouse models of acid beta-glucosidase deficiency: the defect in Gaucher disease. Am J Pathol, 2003. 163(5): p. 2093–101.

82. Furderer, M.L., et al., A Comparative Biochemical and Pathological Evaluation of Brain Samples from Knock-In Murine Models of Gaucher Disease. Int J Mol Sci, 2024. 25(3).

83. Polissidis, A., et al., A double-hit in vivo model of GBA viral microRNA-mediated downregulation and human alpha-synuclein overexpression demonstrates nigrostriatal degeneration. Neurobiol Dis, 2022. 163: p. 105612.

84. Taguchi, Y.V., et al., Glucosylsphingosine Promotes alpha-Synuclein Pathology in Mutant GBA-Associated Parkinson’s Disease. J Neurosci, 2017. 37(40): p. 9617–9631.

85. Kweon, S.H., et al., Gba1 E326K renders motor and non-motor symptoms with pathological alpha-synuclein, tau and glial activation. Brain, 2024. 147(12): p. 4072–4083.

86. Liu, Y., et al., Mice with type 2 and 3 Gaucher disease point mutations generated by a single insertion mutagenesis procedure. Proc Natl Acad Sci U S A, 1998. 95(5): p. 2503–8.

87. Mizukami, H., et al., Systemic inflammation in glucocerebrosidase-deficient mice with minimal glucosylceramide storage. J Clin Invest, 2002. 109(9): p. 1215–21.

88. Mahoney-Crane, C.L., et al., Neuronopathic GBA1L444P Mutation Accelerates Glucosylsphingosine Levels and Formation of Hippocampal Alpha-Synuclein Inclusions. J Neurosci, 2023. 43(3): p. 501–521.

89. Migdalska-Richards, A., et al., Ambroxol effects in glucocerebrosidase and alpha-synuclein transgenic mice. Ann Neurol, 2016. 80(5): p. 766–775.

90. Yun, S.P., et al., alpha-Synuclein accumulation and GBA deficiency due to L444P GBA mutation contributes to MPTP-induced parkinsonism. Mol Neurodegener, 2018. 13(1): p. 1.

91. Frattini, E., et al., Lewy pathology formation in patient-derived GBA1 Parkinson’s disease midbrain organoids. Brain, 2024.

92. Han, T.U., R. Sam, and E. Sidransky, Small Molecule Chaperones for the Treatment of Gaucher Disease and GBA1-Associated Parkinson Disease. Front Cell Dev Biol, 2020. 8: p. 271.

93. Wong, Y.C., et al., Neuronal vulnerability in Parkinson disease: Should the focus be on axons and synaptic terminals? Mov Disord, 2019. 34(10): p. 1406–1422.

94. Vidyadhara, D.J., et al., Synaptic vesicle endocytosis deficits underlie GBA-linked cognitive dysfunction in Parkinson’s disease and Dementia with Lewy bodies. bioRxiv, 2024.

95. Connor-Robson, N., et al., An integrated transcriptomics and proteomics analysis reveals functional endocytic dysregulation caused by mutations in LRRK2. Neurobiol Dis, 2019. 127: p. 512–526.

96. Hunn, B.H.M., et al., Impairment of Macroautophagy in Dopamine Neurons Has Opposing Effects on Parkinsonian Pathology and Behavior. Cell Rep, 2019. 29(4): p. 920–931 e7.

97. Medina-Luque, J., et al., Striatal dopamine neurotransmission is altered in age- and region-specific manner in a Parkinson’s disease transgenic mouse. Sci Rep, 2024. 14(1): p. 164.

98. Rodríguez-Traver, E., et al., GBA1 mutations alter neuronal firing and structure, regulating VGLUT2 and CRYAB in dopamine neurons. bioRxiv, 2024. preprint(DOI: 10.1101/2024.08.05.606574).

99. Cragg, S.J., Variable dopamine release probability and short-term plasticity between functional domains of the primate striatum. J Neurosci, 2003. 23(10): p. 4378–85.

100. Calabresi, P., et al., Alpha-synuclein in Parkinson’s disease and other synucleinopathies: from overt neurodegeneration back to early synaptic dysfunction. Cell Death Dis, 2023. 14(3): p. 176.

101. Aflaki, E., W. Westbroek, and E. Sidransky, The Complicated Relationship between Gaucher Disease and Parkinsonism: Insights from a Rare Disease. Neuron, 2017. 93(4): p. 737–746.

102. Roberts, R.F., R. Wade-Martins, and J. Alegre-Abarrategui, Direct visualization of alpha-synuclein oligomers reveals previously undetected pathology in Parkinson’s disease brain. Brain, 2015. 138(Pt 6): p. 1642–57.

103. Zambon, F., et al., Cellular alpha-synuclein pathology is associated with bioenergetic dysfunction in Parkinson’s iPSC-derived dopamine neurons. Hum Mol Genet, 2019. 28(12): p. 2001–2013.

104. La Vitola, P., et al., Mitochondrial oxidant stress promotes alpha-synuclein aggregation and spreading in mice with mutated glucocerebrosidase. NPJ Parkinsons Dis, 2024. 10(1): p. 233.

105. Fishbein, I., et al., Augmentation of phenotype in a transgenic Parkinson mouse heterozygous for a Gaucher mutation. Brain, 2014. 137(Pt 12): p. 3235–47.

106. Migdalska-Richards, A., et al., L444P Gba1 mutation increases formation and spread of alpha-synuclein deposits in mice injected with mouse alpha-synuclein pre-formed fibrils. PLoS One, 2020. 15(8): p. e0238075.

107. Polinski, N.K., et al., The GBA1 D409V mutation exacerbates synuclein pathology to differing extents in two alpha-synuclein models. Dis Model Mech, 2022. 15(6).

108. Greffard, S., et al., Motor score of the Unified Parkinson Disease Rating Scale as a good predictor of Lewy body-associated neuronal loss in the substantia nigra. Arch Neurol, 2006. 63(4): p. 584–8.

109. Ma, S.Y., et al., Correlation between neuromorphometry in the substantia nigra and clinical features in Parkinson’s disease using disector counts. J Neurol Sci, 1997. 151(1): p. 83–7.

110. Do, J., et al., Behavioral Phenotyping in a Murine Model of GBA1-Associated Parkinson Disease. Int J Mol Sci, 2021. 22(13).

111. Ikuno, M., et al., GBA haploinsufficiency accelerates alpha-synuclein pathology with altered lipid metabolism in a prodromal model of Parkinson’s disease. Hum Mol Genet, 2019. 28(11): p. 1894–1904.

