## Supplemental Figures for "A human *GBA-L444P* transgene drives early and persistent dopamine neurotransmission deficits and alpha-synuclein pathology in a mouse model of early Parkinson’s disease"

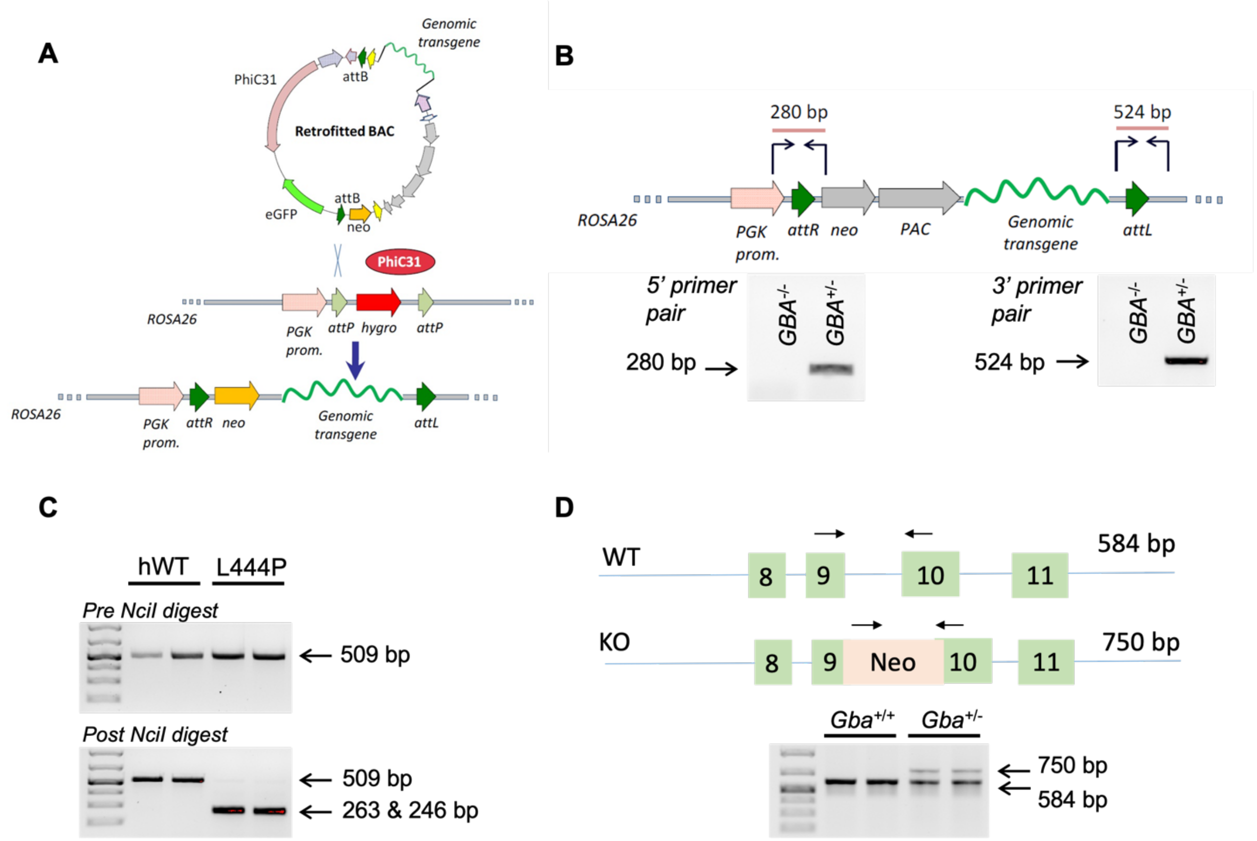


***Supplementary Figure 1: Design of the BAC vector and genotyping for the human GBA transgene and endogenous mouse Gba.*** *(**A**) Schematic of the BAC vector retrofitted with the human GBA transgene. Site specific integration into the Rosa26 locus is achieved through the action of PhiC31 integrase. (**B**) Schematic of the insertion site for the human GBA transgene and representative agarose gel to determine the presence of the human GBA transgenic construct. (**C**) Representative NcII digest reaction to distinguish between the hWT and L444P GBA transgene. The DNA fragment from L444P mice is selectively cleaved by NcII. (**D**) Schematic of the construct containing a neomycin resistance gene inserted between exon 9 and 10 of the Gba gene in Gba*$^{+/-}$ *mice. Representative agarose gels to detect the presence of this construct, thereby determining the zygosity of mouse Gba. bp = base pairs.*

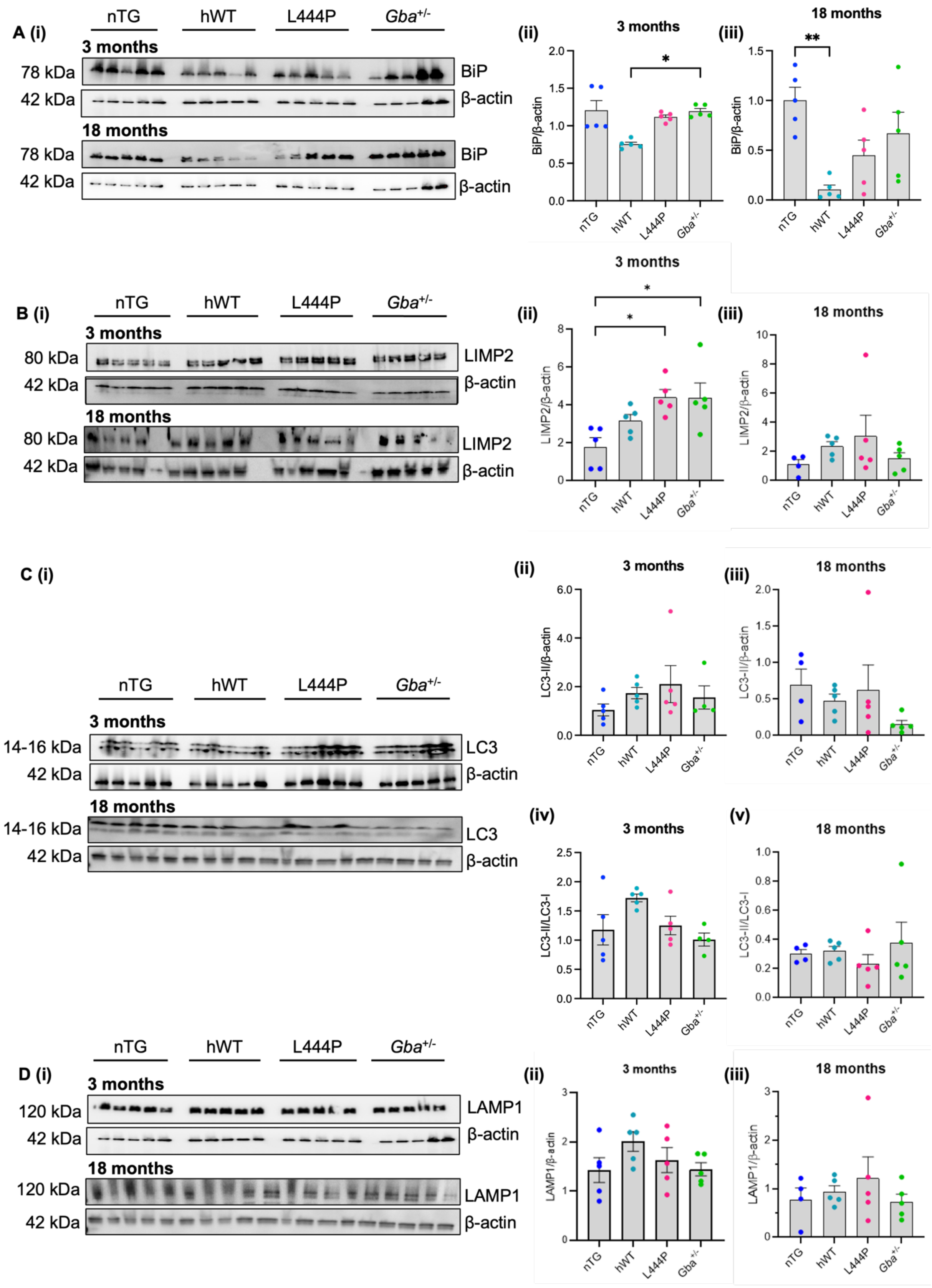
A (i)
nTG
hWT
L444P
Gba+/-
3 months
78 kDa
BiP
42 kDa
β-actin
18 months
(ii)
3 months
BiP/β-actin
(iii)
18 months
B (i)
80 kDa
LIMP2
LIMP2/β-actin
C (i)
14-16 kDa
LC3
LC3-II/β-actin
(iv)
(v)
LC3-II/LC3-I
D (i)
120 kDa
LAMP1
LAMP1/β-actin

**Supplementary Figure 2: Western blots in midbrain tissue from young and aged mice.** Representative western blots (left) and corresponding quantification (right) of LIMP2 (**A**), BiP (**B**), LAMP1 (**C**) and LC3 (**D**) using midbrain tissue samples from 3- and 18-month-old mice. Each protein is normalised to its own β-actin per mouse. (**A(ii), B(iii), D(iv)**) One-way ANOVA with Tukey's post-hoc test: (**A(ii)**) $F_{(3,16)} = 5.529$, $p = 0.0085$, (**B(iii)**) $F_{(3,16)} = 6.484$, $p = 0.0044$, (**D(iv)**) $F_{(3,16)} = 9.084$, $p = 0.0010$; (**A(iii), D(iii)**) Kruskal-Wallis test: (**A(iii)**) Kruskal-Wallis statistic = 3.930, p = 0.2691, (**D(iii)**) Kruskal-Wallis statistic = 6.587, p = 0.0863;(**B(ii), D(ii)**) Kruskal-Wallis test with Dunn's post-hoc test: (**B(ii)**) Kruskal-Wallis statistic = 11.27, p = 0.0103, (**D(ii)**) Kruskal-Wallis statistic = 14.11, p = 0.0028; (**C, D(v)**) One-way ANOVA: (**C(ii)**) $F_{(3,15)} = 0.8769$, $p = 0.4750$ (**C(iii)**) $F_{(3,15)} = 0.6471$, $p = 0.5968$ (**C(iv)** ) $F_{(3,15)} = 03.167$, $p = 0.0553$ , (**D(v)**) $F_{(3,15)} = 0.5265$, $p = 0.6708$. *$p < 0.05$, **$p < 0.01$, ***$p < 0.001$; n = 4-5 mice per genotype (3 months) and n = 4-5 mice per genotype (18 months).

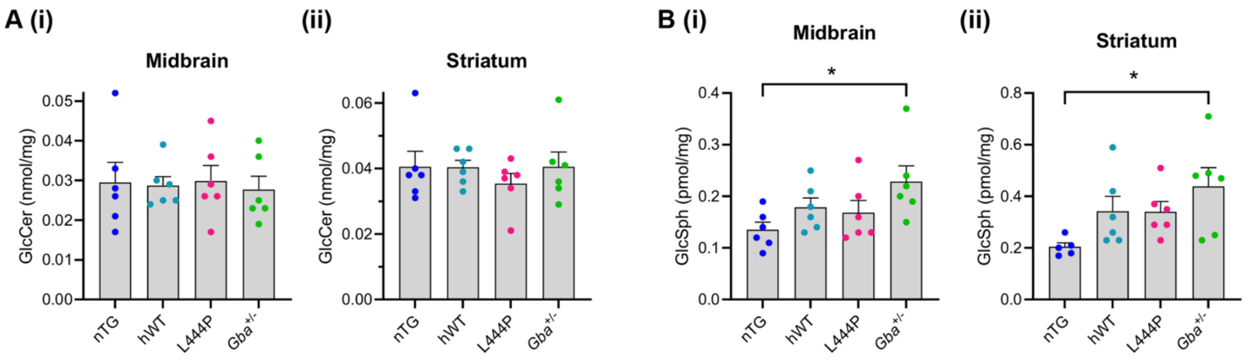


***Supplemental Figure 3: Lipidomics in the midbrain and striatum.*** *Lipidomic analysis of midbrain tissue from 3-month-old mice revealed no changes to glucosylceramide (GlcCer ) levels (**A**) in the midbrain (i) or striatum (ii) across genotypes. However, a significant increase in glucosylsphingosine (GlcSph) (**B**) was observed in Gba$^{+/-}$ mice in both the midbrain (i) and striatum (ii). (**A(i)**) One-way ANOVA: $F_{(3, 20)}$ = 0.06491 and p = 0.9778, (**A(ii)**) Kruskal-Wallis test: Kruskal-Wallis statistic = 1.103 and p = 0.7763, (**B**) One-way ANOVA with Šídák's post-hoc test: (**B(i)**) $F_{(3,20)}$ = 2.878 and p = 0.0616, (**B(ii)**) $F_{(3,19)}$ = 3.070 and p = 0.0527.* *$p < 0.05$; *n = 5-6 mice per genotype.*

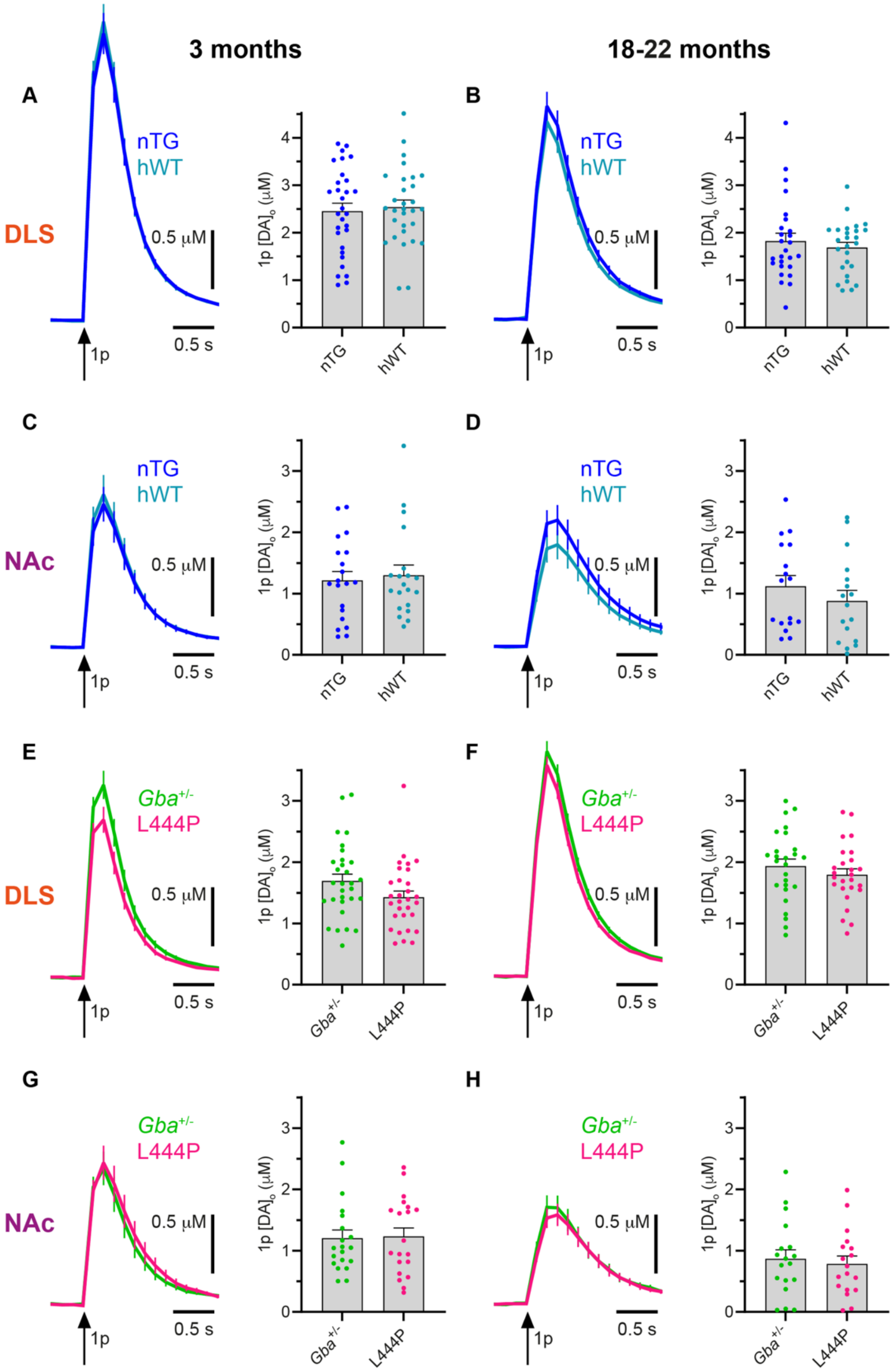


***Suppl. Figure 4: No significant differences in DLS or NAc [DA]$_o$ were observed across pairwise recordings in nTG vs hWT mice and L444P vs Gba$^{+/-}$ mice.** (**A-H**) Mean [DA]$_o$ transients evoked by single pulse electrical stimulations (↑1p) (left) and corresponding peak [DA]$_o$ evoked by the same stimulations (right) in the dorsolateral striatum (DLS) (**A, B, E, F**) and nucleus accumbens core (NAc) (**C, D, G, H**), at 3 months*

*(**A, C, E, G**) and 18-22 months (**B, D, F, H**). nTG and hWT mice had similar $[DA]_o$ in the DLS at 3 months (**A**) and 18-22 months (**B**). (**C-D**) There were also no differences in NAc $[DA]_o$ across nTG and hWT mice at 3 months (**C**) or at 18-22 months (**D**). (**E**) The mean DLS $[DA]_o$ was reduced in L444P mice vs $Gba^{+/-}$ mice at 3 months but this did not reach significance ($p = 0.0606$). (**F**) No significant differences in DLS $[DA]_o$ were observed in L444P vs $Gba^{+/-}$ mice at 18-22 months. (**G-H**) NAc $[DA]_o$ was comparable in L444P and $Gba^{+/-}$ mice at 3 months (**G**) and 18-22 months (**H**). (**A, D, F, H**) Two-tailed paired t-test, (**B-C, E, G**) Wilcoxon matched-pairs signed rank test, all graphs are $p > 0.05$. Each datapoint represents one recording site; (**A, E**) n = 30 recordings from 5 mice per genotype, (**B**) n = 26 recordings from 5 mice per genotype, (**C, G**) n = 20 recordings from 5 mice per genotype, (**D**) n = 17 recordings from 5 mice per genotype, (**F**) n = 26 recordings from 5 mice per genotype, (**H**) n = 18 recordings from 5 mice per genotype.*

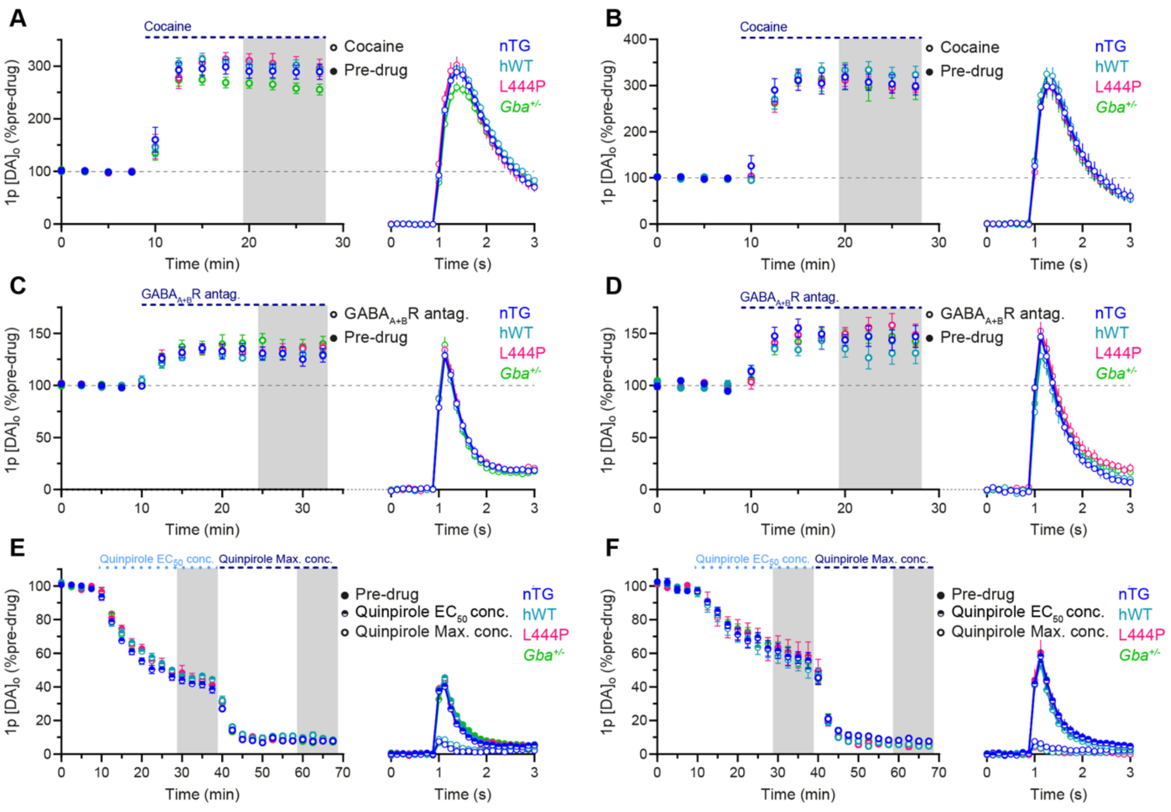


***Suppl. Figure 5: Striatal DAT-mediated DA uptake, $D_2$R-mediated inhibition of DA release and tonic GABAergic inhibition of DA release are not altered in L444P mice.*** *(**A-F**) Left: Peak $[DA]_o$ evoked by single pulse (1p) electrical stimulations at 2.5 min intervals in DLS before (filled datapoints) and during (half-filled or unfilled datapoints) drug wash-on. The period of drug application is indicated by the blue dashed bar. Right: Mean $[DA]_o$ transients during drug wash-on obtained from the four stimulations in the grey shaded areas (left). In all graphs, data are presented normalised within individual genotypes to the mean peak $[DA]_o$ from the four stimulations pre-drug application. Peak $[DA]_o$ was not significantly differentially altered across genotypes at 3-6 months (**A, C, E**) or with age (**B, F**: 15-19 months, **D**: 18-22 months) following application of: (**A-B**) 5 μM cocaine, (**C-D**) $GABA_{A+B}$ receptor antagonists ($GABA_{A+B}$R antag.): 10 μM bicuculine & 4 μM CGP 55845 hydrochloride, (**E-F**) 44 nM ($EC_{50}$) & 1 μM (Max.) quinpirole. This indicates that regulation of striatal DA by the DAT (**A-B**), striatal GABA (**C-D**) and $D_2$Rs (**E-F**) is not altered in L444P mice. (**A-F**) Two-way repeated measures ANOVA: (**A**) interaction: $F_{(7.454,57.15)} = 1.282$, $p = 0.2735$; genotype: $F_{(3, 23)} = 1.943$, $p = 0.1508$, (**B**) interaction: $F_{(7.028,51.54)} = 0.6013$, $p = 0.7526$; genotype: $F_{(3, 22)} = 0.2592$, $p = 0.8539$, (**C**) interaction: $F_{(7.931, 63.45)} = 0.9319$, $p = 0.4963$; genotype: $F_{(3, 24)} = 0.4412$, $p = 0.7257$, (**D**) interaction: $F_{(8.122, 46.03)} = 1.205$, $p = 0.3171$; genotype: $F_{(3, 17)} = 0.8497$, $p = 0.4858$, (**E**) interaction: $F_{(15.33,102.2)} = 0.9217$, $p = 0.5443$; genotype: $F_{(3, 20)} = 1.311$, $p = 0.2986$, (**F**) interaction: $F_{(5.358,$*

*$_{37.51)}$ = 0.4602, p = 0.8148; genotype: $F_{(3, 21)}$ = 0.2113, p = 0.8874. (**A**) n = 5-8 slices from 5-8 mice per genotype, (**B**) n = 5-7 slices from 5 mice per genotype, (**C**) n = 6-8 slices from 6-8 mice per genotype, (**D**) n = 4-7 slices from 4-5 mice per genotype, (**E**) n = 5-7 slices from 5-7 mice per genotype, (**F**) n = 5-8 slices from 4-6 mice per genotype.*

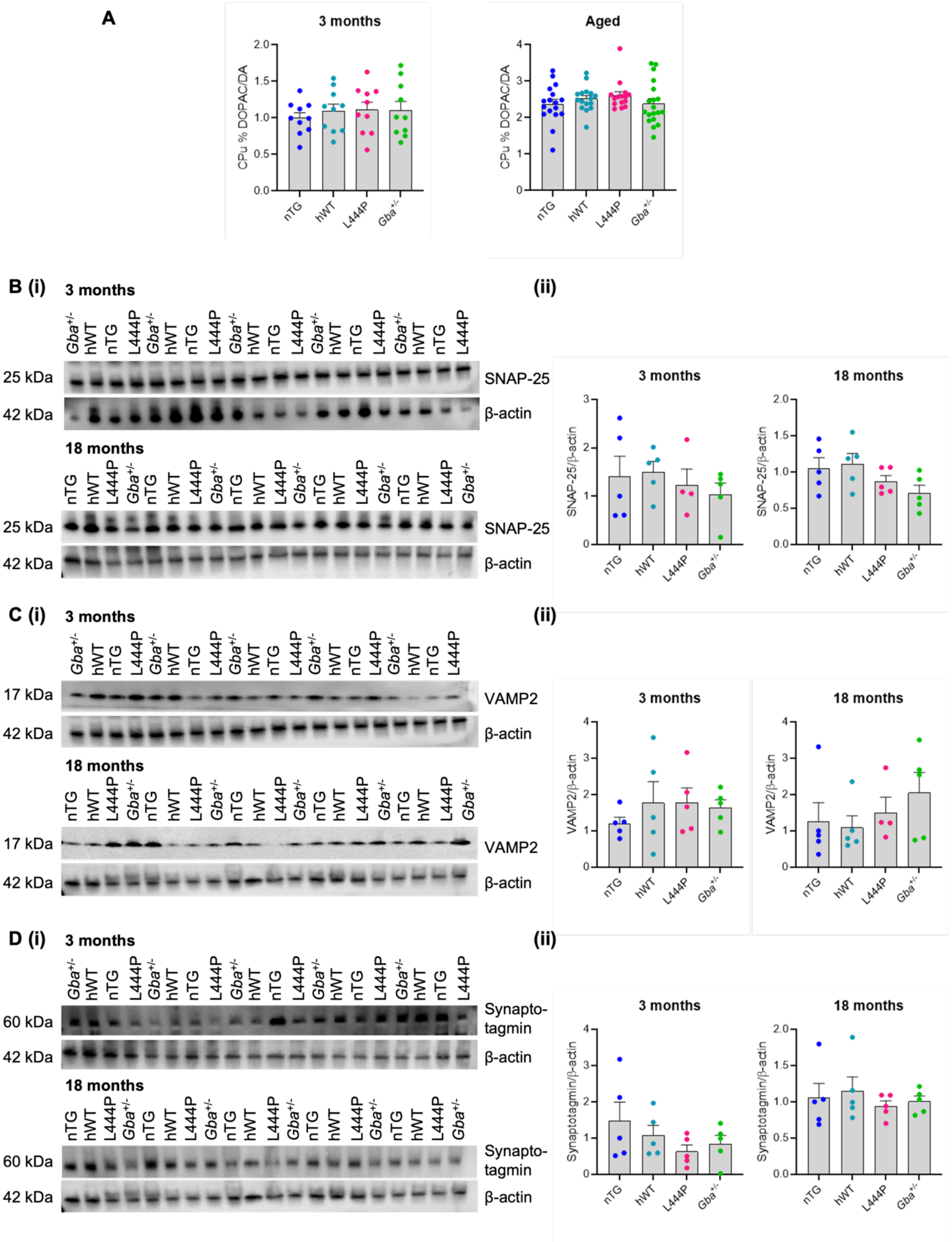


***Suppl. Figure 6: The striatal DOPAC:DA ratio and components of the synaptic release machinery were comparable across genotypes.*** *(**A**) No changes to the DOPAC:DA ratio were observed in the CPu of 3-month-old (**i**) or aged (15-22-month-old) (**ii**) mice, indicating that DA turnover was similar across genotypes. (**B-D**) Representative western blots in striatal tissue and corresponding quantification of components of the*

*synaptic release machinery: SNAP-25 (**B**), synaptotagmin (**C**) and VAMP-2 (**D**). No genotype-related differences were observed in 3-month or 18-month-old mice. (**A(i), B, C(ii), D**) One-way ANOVA: all F < 2.208 and all p > 0.1268, (**A(ii), C(iii)**) Kruskal-Wallis test: all Kruskal-Wallis statistics < 4.156 and all p > 0.2451. (**A(i)**) n = 10 mice per genotype, (**A(ii)**) n = 15-19 mice per genotype, (**B-D**) n = 4-5 mice per genotype.*

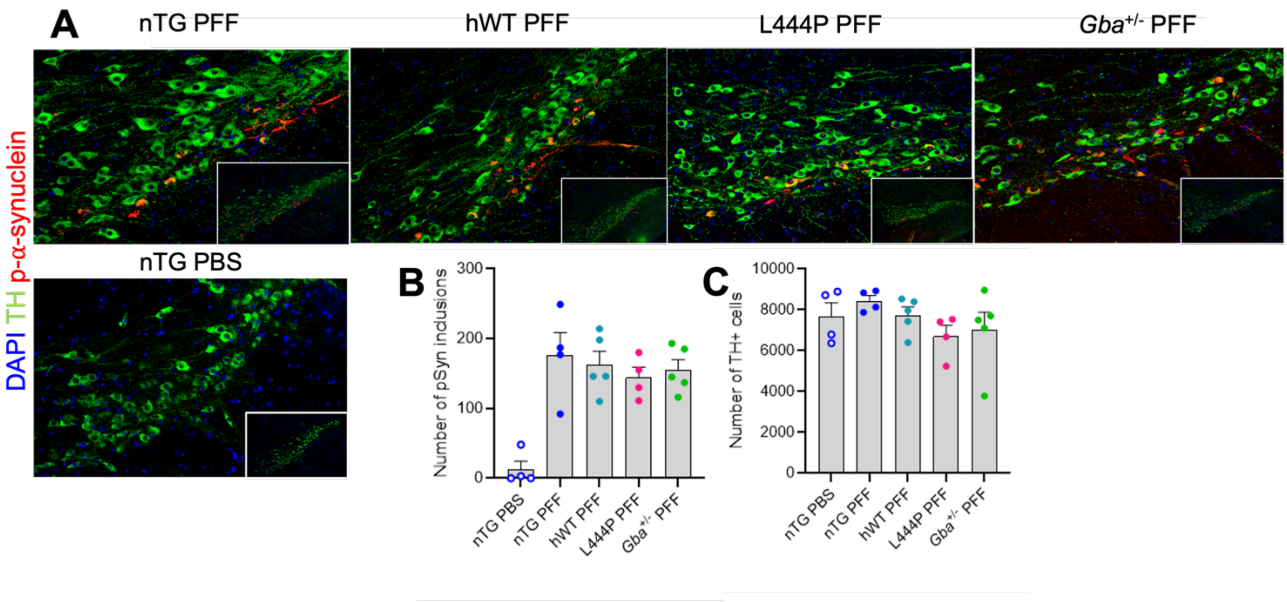


***Suppl. Figure 7:*** **The L444P mutation did not exacerbate PFF-induced α-synuclein pathology in the midbrain.** *(**A**) Representative images of the TH positive cells (green) in the SNpc with p-α-synuclein pathology (red). No significant changes were seen in the number of p-α-synuclein (pSyn) inclusions (**B**), SNpc TH positive cells (**C**). (**B**) Kruskal-Wallis test with Dunn's post-hoc test: Kruskal-Wallis statistic = 10.12, p = 0.0384 (no significance difference by Dunn's post-hoc), (**C, D**) one-way ANOVA: (**C**) $F_{(4,17)}$ = 1.200, p = 0.3468, (**D**) $F_{(4, 18)}$ = 0.6757, p = 0.6175, n = 4-5 mice per genotype.*

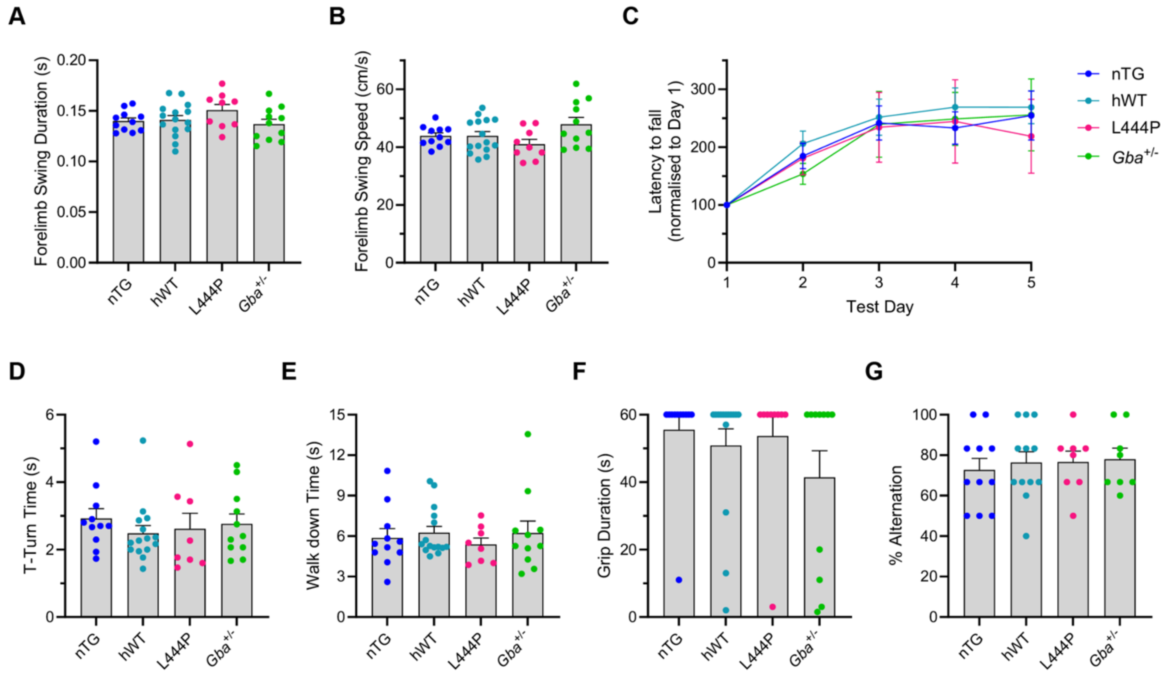


***Suppl. Figure 8: The L444P mutation does not alter forelimb gait dynamics or performance on the rotarod, vertical pole, inverted grid or T-maze assessments.*** *(**A-B**) On the CatWalk XT automated gait analysis system, forelimb swing duration (**A**) and forelimb swing speed (**B**) were similar across genotypes. (**C**) The latency to fall increased to a similar extent in all genotypes upon repeated exposure to the rotarod and no significant genotype-related differences were observed across trial days. Data are presented normalised in each individual mouse to the latency to fall on day one. (**D-E**) The time that mice took to complete a T-turn (**D**) and to walk down (**E**) the vertical pole was comparable across genotypes. (**F**) The time that mice clung to the inverted grid was not significantly different across genotypes; most mice gripped to the underside of the grid for the maximum trial duration of 60 s. (**G**) The spontaneous alternation rate in the T-maze was similar across all genotypes, indicating that spatial working memory is not affected by the L444P mutation. (**A-B, G**) One-way ANOVA: (**A**) $F_{(3,42)} = 1.431$, $p = 0.2472$, (**B**) $F_{(3,42)} = 2.403$, $p = 0.0810$, (**G**) $F_{(3,35)} = 0.1650$, $p = 0.9192$, (**C**) two-way repeated measures ANOVA: interaction $F_{(5.609,78.53)} = 0.2748$, $p = 0.9397$; genotype $F_{(3,42)} = 0.1401$, $p = 0.9354$. (**D-F**) Kruskal-Wallis test: (**D**) Kruskal-Wallis statistic = 2.309, $p = 0.5108$ (**E**) Kruskal-Wallis statistic = 0.8287, $p = 0.8426$ (**F**) Kruskal-Wallis statistic = 3.181, $p = 0.3645$. (**A-C, F**) n = 9-15 mice per genotype, (**D-E**) n = 8-15 mice per genotype, (**G**) n = 8-12 mice per genotype.*
